# Synergistic effects of commensals and phage predation in combating pathogen infections: a shrimp model

**DOI:** 10.1101/2025.03.19.644257

**Authors:** Ling Chen, Zhipeng Huang, Chenli Liu, Mathias Middelboe, Yingfei Ma

## Abstract

Colonization resistance is a fundamental mechanism by which microbiomes suppress pathogen invasion, yet the factors governing its efficacy remain unclear. By constructing a defined microbial consortium and employing *in vivo* models, we provide evidence supporting that phage-mediated pathogen suppression interacts with competitive dynamics within microbial communities to enhance colonization resistance. While individual species conferred limited protection, combining key taxa with phage treatment extensively improved pathogen exclusion. Phage-induced pathogen killing amplified nutrient competition among commensals, however, the timing intervention emerged as a crucial determinant of efficacy. Guided by these ecological principles, we formulated a minimal, synergistic consortium that robustly enhanced colonization resistance *in vivo*. These findings advance our understanding of microbiome-driven pathogen resistance and provide a strategic framework for designing protective microbial consortia with potential applications in human and animal health.

## Introduction

Deciphering the structure and functions of microbiomes, along with their interactions with the host, has increasingly highlighted their crucial roles in modulating the host immune system and maintaining homeostasis(*1*). One critical aspect of this meta-organism immunity is their ability to defend against pathogen colonization, a phenomenon known as colonization resistance(*2*). Colonization resistance involves complex mechanisms, including metabolic competition(*3*), niche occupation(*4*), and the production of antimicrobial agents(*5*). In contrast, dysbiosis refers to a shift from a balanced, diverse microbiome to a state that is more susceptible to pathogen invasion, significantly impacting both microbiome function and host fitness(*6*). Pathogens can actively remodel the microbiota to their advantage, further exacerbating dysbiosis and facilitating infection(*7*, *8*).

Microbiota manipulation, particularly through microbial consortia therapy, has emerged as a promising strategy to restore balance after dysbiosis and combating disease(*9*). In particular, the development of defined microbial consortia, using either engineered or natural commensals, has provided key insights into the role of specific commensals in reinforcing colonization resistance(*10*). However, the adaptive capacity of pathogens enables them to circumvent microbiome-based defenses, posing a major challenge for microbiota-targeted therapies(*11*, *12*). Bacteriophages (phages) are the most numerous biological entities cross earth’s biospheres, playing a pivotal role in shaping microbial communities and controlling bacterial populations(*13*). Their remarkable host specificity makes them powerful tools for pathogen suppression, with increasing interest in their potential to synergize with microbiome modulation for enhanced disease resistance(*14*).

Among marine pathogens, *Vibrio* species are ubiquitous in aquatic ecosystems, with some strains serving as commensals, while others cause devastating infections in marine animals and humans even(*15*, *16*). *Vibrio parahaemolyticus*, particularly strains carrying the *pirAB* toxin genes, is the primary etiological agent of acute hepatopancreatic neurosis disease (AHPND) in shrimp, a condition associated with severe hepatopancreatic damage and mass mortality(*17*). Advance in next-generation sequencing have revealed significant microbiome shifts during *Vibrio* infections, with diseased shrimp consistently exhibiting an overrepresentation of *Vibrio* species in their gut communities(*18*). While *Vibrio*-induced dysbiosis is well documented, the role of commensal *Vibrio* species in defending against their pathogenic counterparts remains poorly understood. Given the ecological complexity of *Vibrio* populations within host-associated microbiomes, unraveling the protective functions of non-pathogenic strains could provide novel insights into microbiota-driven pathogen resistance.

Building upon these insights, we hypothesized that a synergistic approach integrating commensal *Vibrio* species with phages application could enhance pathogen suppression through microbiome-mediated resistance and targeted phage predation. To test this, we established an experimental model simulating pathogen invasion, microbiome colonization resistance, and phage predation in shrimp. Our findings demonstrate that combining a commensal microbial consortium with a *Vibrio*-specific lytic phage provides superior pathogen suppression compared to either strategy alone. Furthermore, we elucidate the ecological mechanisms underlying microbiome-mediated defense, emphasizing the role of ecological diversity and priority effects in shaping pathogen dynamics. These results highlight the potential of microbiome-phage synergy for pathogen control, potentially with broader implications for microbiome-based disease management strategies in host-associated ecosystems.

## Results

### Commensal intestinal bacteria can protect shrimps from the pathogenic *Vibrio* infection

To investigates the role of commensal bacteria in protecting shrimp from pathogenic *Vibrio* infections, we first characterized the dominant phyla in the healthy shrimp gut microbiome. Based on V3-V4 of 16S rRNA gene amplicon sequencing from our previous study(*19*), the predominant phyla included Proteobacteria, Bacteroidetes, Firmicutes, and Actinobacteria. Given their prevalence, we selected representative strains from these groups to construct a bacterial consortium, termed “**Com12**”.

Com12 comprises of twelve distinct commensal strains isolated from the shrimp intestine, and taxonomically classified via whole-genome sequencing (see **Methods**). The consortium includes species from the major phyla: Proteobacteria (e.g., *Halocynthiibacter* sp. [Halo], *Ruegeria* sp. [Rueg], *Shewanella* sp. [Shew], *Pseudoalteromonas* sp. [Pseu], *Psychrobacter* sp. [Psyc]; Bacteroidetes (e.g., *Tenacibaculum* sp. [Tena], *Algoriphagus* sp. [Algo], *Gaetbulibacter* sp. [Gaet]); Firmicutes (e.g., *Exiguobacterium* sp. [Exig], *Planococcus* sp. [Plan]); and Actinobacteria (e.g., *Microbacterium* sp. [Micr], *Demequina* sp. [Deme]) (**Extended Data Table 1**).

To examine interactions between commensal and pathogenic bacteria, we conducted *in vitro* co-culture experiments with Com12 consortium and two *Vibrio* strains: one pathogenic (*Vibrio parahaemolyticus* strain VP6) and one putatively beneficial (*Vibrio alginolytic* strain VA3). Initially, taxonomic characterization of these two *Vibrio* strains was performed via whole-genome sequencing, and phenotypic classification was based on the presence of virulence-associated genes and mortality assays. Genome annotation revealed that VP6 carries the *pirAB* toxin gene, whereas VA3 lacks this virulence determinant, a finding further confirmed by genetic analysis (**Extended Data Fig. 1**). Infection experiments demonstrated that VP6 induced significant mortality and vibriosis-specific symptoms, yet shrimp exposed to VA3 exhibited no detectable difference in survival relative to the control group (**Fig. 1a, Extended Data Fig. 2**). Co-culture experiments further revealed that both *Vibrio* strains significantly altered the structure of the Com12 consortium, with each strain becoming dominant within the microbial community (**Extended Data Fig. 3**). Notably, VA3 exhibited a distinct growth advantage over VP6 in Com12, suggesting that it may competitively inhibit VP6 colonization.

**Fig. 1.**
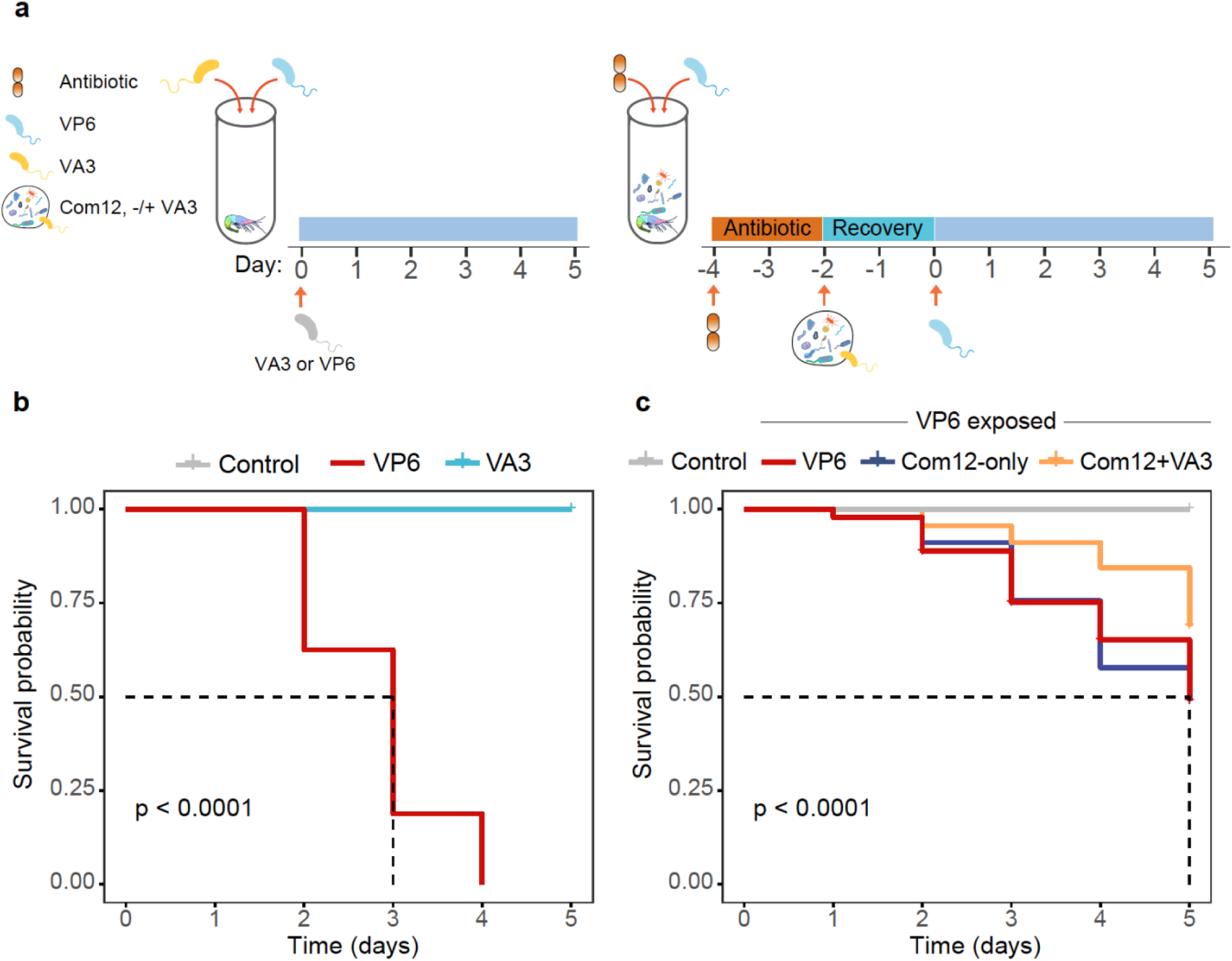
Characterization of commensal and pathogenic strains in shrimp. **a**, Overview of the *in vivo* assays. In the invasion assay (**left**), shrimp was housed according to standard procedures use in shrimp farming before the addition of the pathogen VP6 or the symbiont VA3. In the alternative invasion assay (**right**), shrimp were pre-exposed to a combination of antibiotics (see **Methods**) and after two days, were randomly divided into groups and bathed in seawater containing different bacterial strains. The Com12 consortium or individual strains were pre-grown in marine broth medium (2216E) prior to the addition of the pathogen (VP6). **b**, Survival rate curves of shrimp exposed to each vibrio strains (VP6 and VA3). Fresh bacterial cultures, adjusted to 5×10^6^ CFU/mL, were added to the shrimp house environment. A control group of shrimp without any treatment was included for comparison. Median survival was indicated by black dotted lines in panel. Statistical significances were calculated between the groups using the log-rank test. Statistical results are given as exact *P* values or indicated *P* < 0.0001. **c**, Survival rate curves of antibiotic-treated shrimp. Groups include: control (shrimp treated with antibiotics), VP6 group (antibiotic-treated shrimp exposed to VP6), VP6+Com12 group (antibiotic-treated shrimp exposed to VP6 and the Com12 consortium), and VP6+Com12+VA3 group (antibiotic-treated shrimp exposed to VP6, the Com12 consortium and VA3). Median survival was indicated by black dotted lines in panel. Statistical significances were calculated between the groups using the log-rank test. Statistical results are given as exact *P* values or indicated *P* < 0.0001.

To further assess the protective role of VA3, we pretreated shrimp with antibiotics to minimize the native microbiota and subsequently recolonized some individuals with the Com12 consortium, -/+ VA3, at a concentration of 5×10^6^ CFU/mL. All shrimp were then challenged with VP6 (5×10^6^ CFU/mL). By day 5, the VP6-only group exhibited 100% mortality (**Fig. 1b**). In contrast, both the Com12+VA3 and Com12-only groups showed increased survival, with the former achieving a significantly higher survival rate (69 %) compared to the Com12-only group (49 %, *P* < 0.05; **Fig. 1b)**. These results highlight the role of VA3 in enhancing the Com12-based resistance to VP6 infection. However, further investigations are needed to determine whether VA3 alone is sufficient to confer protection in the absence of Com12−an open question that warrants deeper exploration in future studies.

In addition, to explore the potential of phage therapy in augmenting microbiota-mediated colonization resistance, we isolated an obligate lytic myovirus, VP6phageC, using VP6 as the host (**Extended Data Fig. 4a,b**). The host range of VP6phageC was assessed against each monospecies within the Com12 consortium, including VA3, using phage infection assays(*20*). As expected, VP6phageC failed to form plaques on any Com12 members (**Extended Data Table 2)**, confirming its strict specificity for VP6 and the absence of cross-infection within the consortium. This specificity makes VP6phageC as a promising candidate for investigating whether a combined approach−leveraging phage predation alongside microbiota modulation, can further enhance shrimp defenses against VP6 colonization.

### Commensal bacteria and phage as dual mechanisms of preventing *Vibrio* pathogen invasion

To investigate the interplay between commensal bacterial and phage in resisting *Vibrio* pathogen invasion, we employed Com12 as a model system to simulate pathogen invasion and evaluate the combined effects of commensal-derived colonization resistance and phage predation. These experiments were performed on a 96-well microplate platform, using time-resolved measurements of mono-species to track the dynamic changes over 48-hour period (**Fig. 2a**, see **Methods**).

**Fig. 2.**
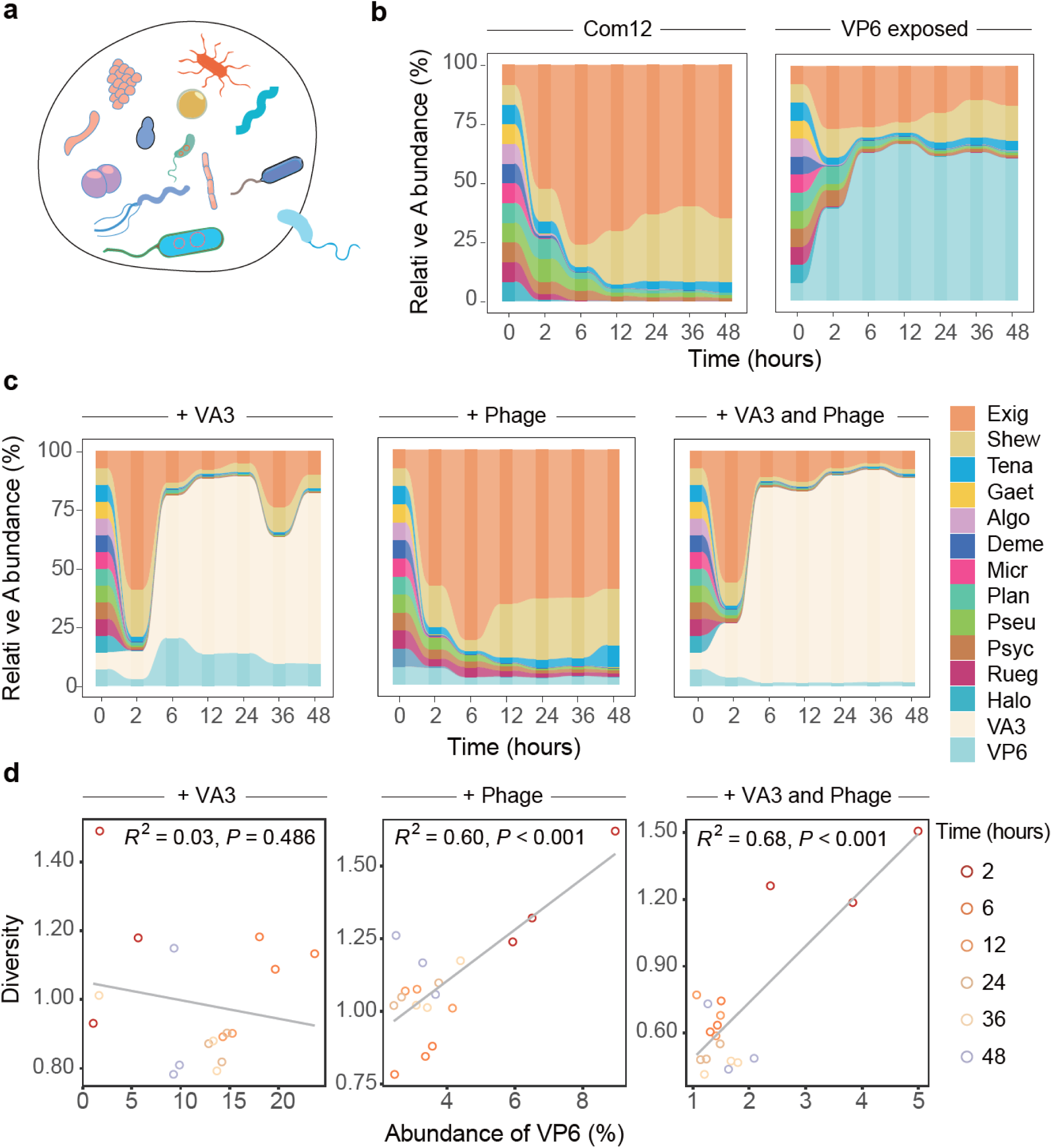
Temporal changes in the composition of the Com12 consortium under different treatments. **a**, Schematic of the *in vitro* experimental design to stimulate pathogen invasion. The Com12 consortium was constructed using 12 distinct strains (at the species level), mixed in approximately equal initial proportions based on OD600 values. Relative abundance was assessed via multiplexed 16S rRNA gene sequencing of the V4 region (see **Methods**). In invasion assays, a 1:1 ratio of VP6 culture was added into the Com12 consortium. Samples were collected at the following time points: 0 h, 2 h, 6 h, 12 h, 24 h, 36 h, and 48 h for sequencing, unless otherwise noted. **b**, Temporal changes in relative abundance of microbial species in the Com12 consortium, **unexposed** (**left**) or **exposed** (**right**) to VP6. The stacked area plot shows the relative abundance of each species over time (y-axis: relative abundance, x-axis: time in hours). **c**, Temporal changes in the composition of the Com12 consortium exposed to pathogen VP6 and additional treatments: **+VA3** (**left**), **+Phage** (**center**), and **+VA3 and Phage** (**right**). Each plot shows the relative abundance of community members within Com12 over time, highlighting the effect of VA3 and phage treatments on community dynamics, individually or together. **d**, Correlation between the diversity index (Shannon) and VP6 abundance at different time points (2, 6, 12, 24, 36, and 48 hours). The plots show the relationship between the dynamics of microbial diversity within the Com12 consortium and VP6 abundance, with R-square and *P* values provided for each treatment (**+VA3** (**left**), **+Phage** (**center**), and **+VA3 and Phage** (**right**)). Circles of varying colors correspond to data from different time points, with solid gray lines reflecting correlation (linear regression, see **Methods**).

Commensal-mediated resistance was evaluated by co-culturing Com12 and Com12+VA3 with VP6 and monitoring strain abundance over time. VP6 rapidly dominated Com12, reaching 70 % relative abundance within 6 hours before stabilizing (**Fig. 2b**). However, the presence of VA3 significantly restricted VP6 expansion, limiting its abundance to 15 %. Phage addition further suppressed VP6 to 5 %, and the combination of VA3 and phage nearly eliminated it, reducing its abundance to just 1 % (**Fig. 2c)**.

Further analysis of the co-culture dynamics revealed notable changes in bacterial abundance and diversity. The introduction of either VA3 or phage, as well as their combination, led to a marked decrease in VP6 abundance, while the abundance of other strains within the Com12 consortium increased (**Fig. 3b**). A correlation analysis between VP6 relative abundance and Com12 diversity revealed that VA3 alone had no significant effects on the relationship between VP6 abundance and Com12 diversity (R-square = 0.03, F-statistics = 0.51, *P* = 0.486). In contrast, phage treatment showed a positive correlation with diversity (R-square = 0.60, F-statistics = 24.30, *P* < 0.001). When VA3 and phage were combined, while the correlation between VP6 abundance and diversity was reduced, the positive relationship between diversity and pathogen suppression was maintained (R-square = 0.68, F-statistics = 33.30, *P* < 0.001) (**Fig. 2d**).

**Fig. 3.**
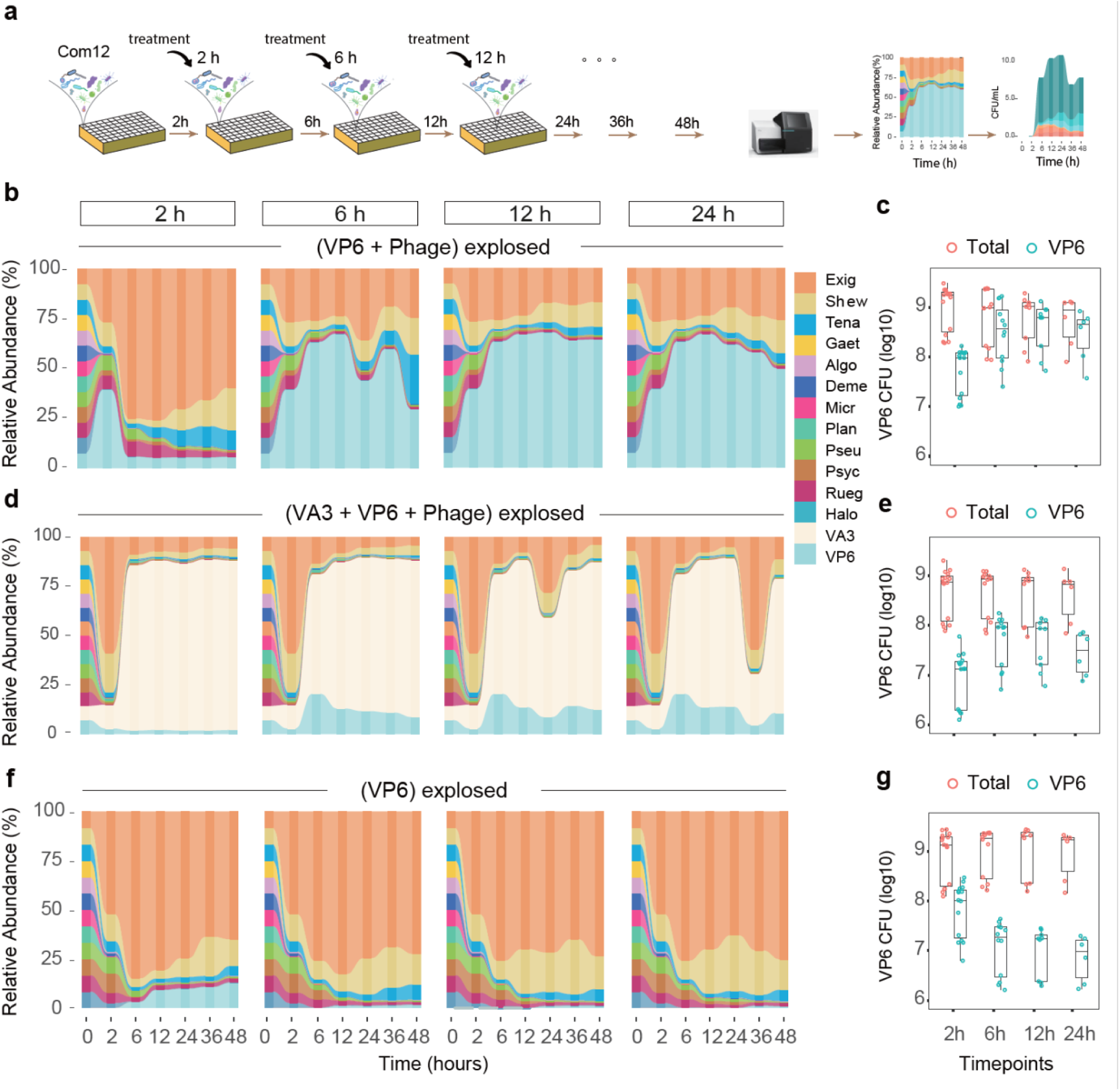
Timing-dependent effects of different treatments on the colonization resistance of the Com12 consortium upon exposure to pathogen VP6. **a**, Schematic of the *in vitro* experimental design modeling pathogen invasion and commensal microbiota colonization resistance. Phage was introduced at specific time points by adding a phage lysate (1:1 ratio, MOI = 1) to the Com12 consortium. Samples were collected at 0 h, 2 h, 6 h, 12 h, 24 h, 36 h, and 48 h for sequencing. Quantification was performed using *E.coli* MG1655 (3.65×10^6^ CFU) as an interior marker (see **Methods**). **b**, Temporal changes in the composition of the Com12 consortium following pathogen VP6 exposure, with phage introduced at specific time points. The time of phage addition is indicated above each subplot. Similar annotations for ***d*** and ***f***. **c,** Dynamics of bacterial and VP6 concentrations following phage treatment at the indicated time points. Data points represent total bacterial concentrations (salmon color) and VP6 concentration (cyan color), collected at 2 h, 6 h, 12 h, 24 h, 36 h and 48 h. **d**, Temporal changes in the composition of the Com12 consortium following pathogen VP6 exposure, with phage added at specific time points, preceded by VA3 treatment of the consortium. **e,** Dynamics of bacterial and VP6 concentrations with phage treatment at the indicated time points, after VA3 was initially added to the Com12 consortium. Data points represent total bacterial concentrations (orange) and VP6 concentration (sky blue), collected at 2 h, 6 h, 12 h, 24 h, 36 h and 48 h. **f**, Temporal changes in the composition of the Com12 consortium following VP6 exposure at specific time points. **g**, Dynamics of bacterial and VP6 concentrations with VP6 introduced at the indicated time points. Data points represent total bacterial concentrations (orange) and VP6 concentration (sky blue), collected at 2 h, 6 h, 12 h, 24 h, 36 h and 48 h.

These results demonstrate that while Com12 alone impose a threshold on VP6 colonization, VA3 and phage act as potent inhibitors, with their combined application synergistically enhance colonization resistance. Importantly, phage contributed to increase microbial diversity, whereas VA3 appears to influence pathogen abundance without directly altering diversity, suggesting complementary mechanisms in pathogen exclusion. Together, these results underscore the potential of integrating commensals and phages as strategy to fortify maintaining microbiome stability and prevent pathogen invasion.

### Timing and synergy in colonization resistance against pathogen invasion

In our study of the Com12 consortium, we identified a priority effect that influenced pathogen invasion, particularly when VP6 was introduced at different stages of Com12 growth. To evaluate how phage treatment could restore the consortium’s resistance following VP6 invasion, we reconducted co-culture experiments where Com12 was exposed to VP6, and VP6phageC (MOI = 1) was introduced at various time points (see **Methods**). Our findings indicated that the timing of phage introduction significantly affected its efficacy (**Fig. 3a)**. More specifically, when VP6 was co-cultured with Com12 for 6 hours or more before the addition of phage, the suppressive effect of the phage was notably diminished, with VP6 relative abundance surged to 70 % of the community, suggesting that phage-mediated suppression was less effective after this time window (**Fig. 3b**). A comparison of absolute VP6 concentrations using copy numbers (unless otherwise specified) in the consortium further corroborated this observation, showing a marginal but not statistically significant reduction in VP6 when phage was added after 6 hours (*P* > 0.05) (**Fig. 3c**). These results suggested that the efficacy of phage treatment is compromised once VP6 has had a chance to establish itself within the consortium for an extended period.

Next, we explored the potential for combining certain commensal species with phage-specific predation to further bolster colonization resistance. Specifically, we assessed the effects of introducing commensal VA3, alongside the lytic phage VP6phageC. In these experiments, VA3 was introduced into Com12 consortium following VP6 invasion, with concurrent VP6phageC treatment. Monitoring of the consortium dynamics revealed significant inhibition of VP6 by this combined treatment. While introducing VA3 alone, showed a significant reduced the relative abundance of VP6 to 15 % and its biomass to 5×10^7^ CFU/mL within 48 hours (**Fig. 2c**), the combined treatment of VA3 VP6phageC further amplified the inhibitory effect. In this dual intervention, VP6 proliferation was almost entirely eradicated, with its relative abundance dropping below 1% and biomass reduced to less than 1×10^6^ CFU/mL (**Fig. 3d,e**). Even when phage was introduced after a 6-hour delay, both VP6 relative abundance and biomass remained significantly lower compared to treatments using either VA3 or phage alone (*P* < 0.001) (**Fig. 3d,e)** These results underscore that while phage-mediated pathogen suppression is highly time-dependent, its effectiveness can be synergistically enhanced when combined with commensal bacteria such as VA3, which together provide robust colonization resistance and protect the microbiota from pathogen invasion.

To further investigate whether the Com12 consortium itself possesses intrinsic, self-regulated resistance to pathogen invasion, we conducted an additional experiment by introducing VP6 at various stages of Com12 growth. Relative abundance analyses revealed a notable difference in outcomes depending on the timing of VP6 introduction. When VP6 was co-cultured with Com12 form the start (0 hour), it quickly dominated the consortium as reflected in its high relative abundance and biomass (**Fig. 2b, upper right**). In contrast, when VP6 was introduced after 2 hours of Com12 growth, its proliferation was irreversibly inhibited, with VP6’s absolute concentration falling to less than 1×10^5^ CFU/mL, and its relative abundance dropped below 1 % (**Fig. 3f,g**).

Overall, these findings underscore a critical, timing-dependent trait of colonization resistance within the consortium, suggesting that early establishment of the commensal consortium provided a robust barrier against pathogen invasion., underscoring the importance of microbial community maturation. Conversely, allowing the pathogen to establish dominance before the consortium has fully matured significantly undermined the consortium’s resistance mechanisms.

### Interactions and functional similarity evaluation pinpoints certain species in Com12 in conferring resistance the pathogenic *Vibrio* invasion

To explore the potential mechanisms underlying the observed colonization resistance in above co-culturing experiments (**Fig. 2-3**), we investigated pairwise interactions among members of the consortium Com12, including *V. parahaemolyticus* (VP6) and *V. alginolyticus* (VA3), using a conditional coculturing approach(*21*). In this experiment, each species was grown in cell-free spend media collected from other species, supplemented with 60 % full-nutrient Marine Broth (MB2216). This experimental design eliminated direct cell-cell contact as a potential mechanism for colonization resistance, allowing us to focus solely on metabolic-mediated interactions.

Analysis of the growth rate and maximum biomass (OD600) of each species grown in spend media from others versus self-derived media revealed a negative correlation, though the absolute correlation index less than 1 (Estimate = -0.51, R-square = 0.12, F-statistic = 13.13, *P* = 2.57e-06) (**Fig. 4a, left)**. This suggests that most interspecies interaction were inhibitory, albeit the extent of inhibition primarily affected total biomass rather than growth rate. When examining the effects of other species on VP6, we observed a similar inhibitory trend, but with a stronger negative correlation (Estimate = -1.31, R-square = 0.54, F-statistic = 23.60, *P* = 0.004) (**Fig. 4a, right**). Ranking the effects of others on VP6 showed that the inhibitory effects were predominantly self-driven or only slightly mediated by VA3, particularly when excluding biomass (**Fig. 4a, right**). This indicates that nutrient overlap with commensals in Com12 had minimal impact on VP6 growth individually, suggesting that other metabolic or ecological factors may be contributing to VP6 suppression in the consortium.

**Fig. 4.**
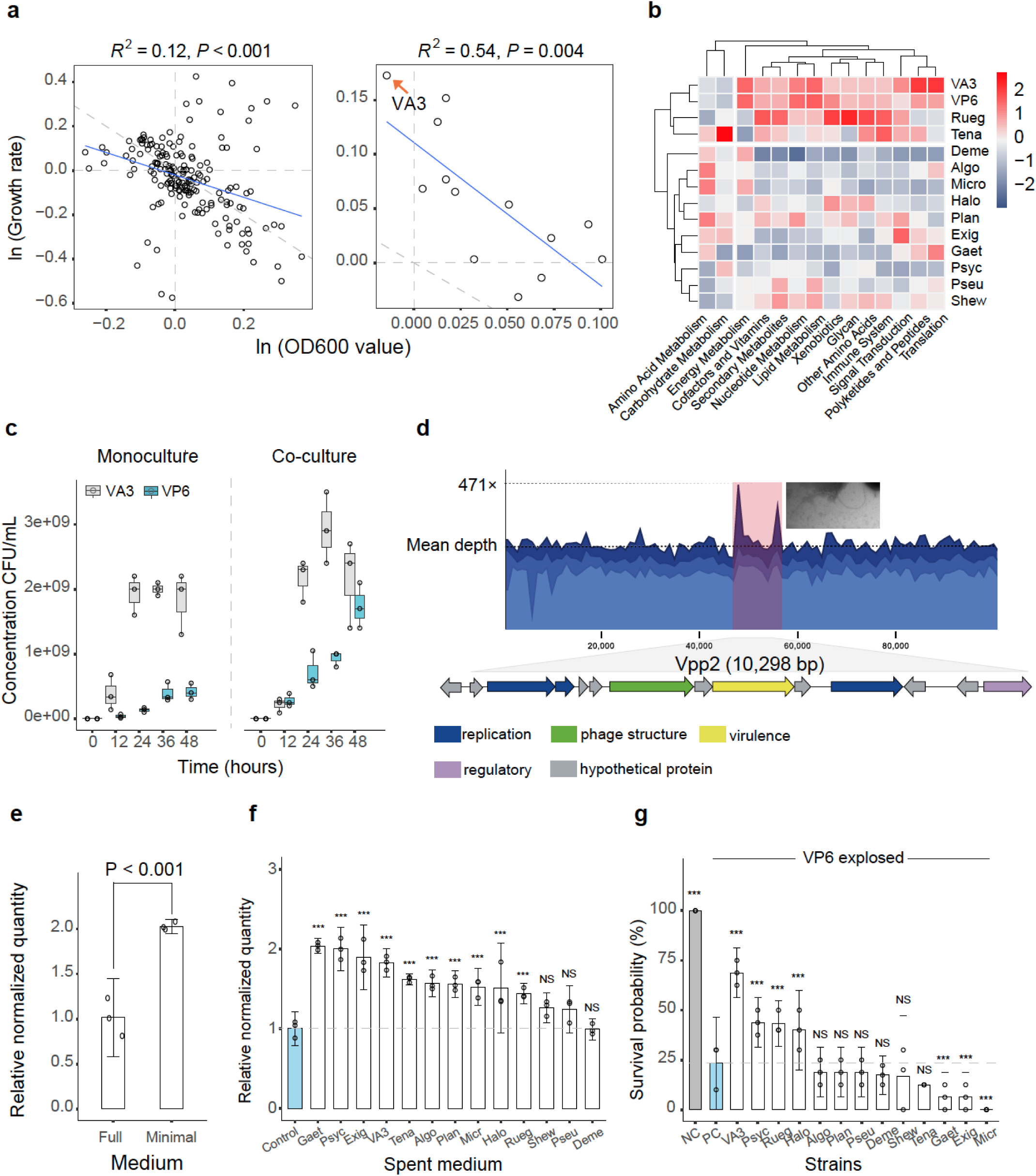
Characterization of the interactions between commensals, VP6 and VA3. **a**, Scatterplots showing the correlation between maximum biomass and growth rate among commensals, VP6 and VA3. Growth was assessed by comparing each strain’s performance when grown in the spend medium of the other species versus its own. More specifically, *a* species grown in the spend medium of *b* exhibited growth rate (Rs) and maximum biomass Kms, while growth in its own spend medium resulted in growth rate (Ro) and maximum biomass (Kmo). Data were plotted as [ln(Km0/Km0] on the x-axis and [ln (Ro/Rs)] on the y-axis. **Left**: Interactions among commensals, VP6 and VA3 (two-sided test, y = -0.51x+b). **Right**: Effect of each commensal within the Com12 consortium and VA3 on VP6 growth (two-sided test, y = -1.3x+b). The solid blue lines show regression lines, and the slash gray dashed lines represent 1:1 reference lines. Linear correlations between the maximum biomass ratio [ln (Kmo/Kms)] and growth rate ratio [ln (Ro/Rs)] are shown with corresponding *P* values. **b**, Heatmap showing gene-level overlap in the Com12 consortium, including VA3 and pathogen VP6. Gene annotations were clustered at the family level of the Pathway family and scaled according to the total gene number within each family. **c**, Growth competition between VA3 and VP6 in co-culture. Overnight cultures of VA3 and VP6 were adjusted to the same OD600 in fresh medium and transferred into a new culture at equivalent concentrations. The concentration of each strain was quantified by plating with serial dilutions on selective agar plates, with distinct morphologies (size difference) allowing for separate enumeration of VA3 and VP6. **d**, Relative abundance of VP6 reads when cultured in minimal medium (SM condition). The genome region of the prophage Vpp2 integrated into VP6 genome is annotated and colored based on the detected molecular functions of individual gene products. Transmission electron microscopy (TEM) images of prophage Vpp2 excision reveal a filamentous morphology typical for inoviridae prophages. Scale bar (white line), 200nm. **e-f**, Relative quantities of prophage Vpp2 DNA when cultured in SM condition medium (***e***) or bacterial condition medium (***f***). Full refers to VP6 cultured in marine broth. In panel *f*, dashed line indicates a reference value of y = 1. In panels ***e*** and ***f***, a two-sided Student’s *t*-test was used. Statistical results are given as exact *P* values in brackets in the graphs or indicated using the following: NS, not significant (*P* > 0.05, unless specified otherwise in the figure), \*\*\**P* < 0.0001. **g**, Impact of Com12 consortium strains and VA3 on shrimp survival after exposure to VP6. Antibiotic-treated shrimp were immersed in cultures of each strain (5×10^6^ CFU/mL) for 2 days, followed by exposure to VP6. Survival rates were calculated based on the number of living shrimp on day 5. NC (negative control) denotes shrimp treated with antibiotics only, and PC (positive control) represents shrimp exposed to VP6 after antibiotic treatment. The dash line represents survival rates of shrimp in the positive group. Statistical significance was assessed using a two-sided Student’s *t*-test with pair comparisons to VP6, with results indicated using the following: NS, not significant (*P* > 0.05, unless specified otherwise in the figure), \**P* < 0.01, \*\*\**P* < 0.0001.

To further explore the metabolic interactions underlying these observations, we employed genome-scale metabolic modeling to assess the functional similarity between VP6 and each member of Com12, including VA3, based on protein composition overlap. Specifically, we quantified the proportion of protein families carried by VP6 that were also shared in each commensal (see **Methods**). Our results highlight that VA3, Rueg, and Tena as key contributors to protein-family overlap with VP6, suggesting their potential role in shaping VP6’s growth dynamics (**Fig. 4b**). Notably, VA3, belonging to the same bacterial genus as VP6, exhibited the highest degree of overlap, reinforcing its potential for strong competitive interactions.

To experimentally validate the pairwise interaction between VA3 and VP6, we conducted direct growth competition assays, in which a 1:1 mixture of VA3 and VP6 cells was co-cultured in nutrient-rich medium (Marine Broth) for 48 hours, alongside monoculture controls. VA3 displayed robust growth, achieving cell densities comparable to its monoculture controls (*P* = 0.52) (**Fig. 4c**). In contrast, VP6 growth was severely impaired in the presence of VA3, with its cell density significantly reduced by 2-to 8-fold (*P* = 0.004) during the 48-hours competition period (**Fig. 4c**). Furthermore, VA3 consistently outcompeted VP6 in both total biomass (0.97 vs 0.75) and growth rate (0.13 vs 0.10, by hour) (**Fig. 4c, Extended Data Fig. 5**). Together, these results align with the suppression observed in Com12 upon the introduction of VA3 (**Fig. 2,3**), reinforcing the notion that VA3 inhibits VP6 proliferation. The observed suppression appears to be primarily due to nutrient competition, particularly the overlap in nutrient utilization between the two strains. We didn’t go further examining the direct antagonistic effects considering that no significant impact of conditioned media from VA3 on the maximum growth of VP6 when compared with the self-induced effect (**Fig. 4a, left**).

### Prophage dynamics in VP6 and implication for microbial interactions and keystone strain-mediated shrimp protection

A recent study revealed that prophages in *Vibrio* strains are inducible and play critical roles in strain competition within marine environments^30^. Inspired by this, we identified two intact prophages in the genome of *Vibrio* VP6. Under nutrient-limited conditions, one prophage was highly induced, as evidenced by increased read depth in its corresponding genomic region (**Fig. 4d**). Transmission electron microscopy (TEM) of the filtered supernatant confirmed the presence of filamentous phage particles characteristic of Inoviruses, measuring approximately 1,800-2,000 nm in length and 5 nm in width (**Fig. 4d, upper panel**). The genome of this phage, termed Vpp2, closely resembled that of filamentous phages based on its size (10,298 bp) and gene annotation (**Fig. 4d, lower panel**).

To investigate the role of prophage Vpp2 in microbiota interaction, we assessed its induction in conditioned media from 13 donor strains, using media from VP6 as a control. Vpp2 production increased in all conditioned media except that from Deme (**Fig. 4e,f**). When nutrient-deficient SM buffer was used, Vpp2 production also increased, indicating that nutrient limitation is a key factor trigger for its activation (**Fig. 4e**). Co-culturing VP6 with VA3 or Com12 separately revealed continuous induction of Vpp2 during 48 hours of co-culture with VA3, whereas the induction was less pronounced with Com12 (**Extended Data Fig. 6**). Together, these findings indicate that both Com12 and commensal VA3 promote prophage induction in VP6, with VA3 exhibiting the most robust effect. This induction likely involves nutrient competition, which activates a stress response in VP6, and suggests that prophage activation may be part of a broader ecological strategy that influences the growth dynamics of VP6.

The dynamics of the Com12 consortium, -/+VA3, revealed that individual strains contribute variably to colonization resistance against pathogen invasion, with some strains playing more pivotal roles. To evaluate the protective capacity of each commensal, we further assessed shrimp survival following exposure to pathogen VP6 (**Fig. 4g**). Shrimp were pretreated with antibiotics as before (**Fig. 2g**) to minimize the influence of indigenous bacteria and then immersed in cultures of each strain (5×10^6^ CFU/mL) or combinations, including Com12 and VA3, prior to VP6 exposure. With this assay system, we could rank the strains based their abilities to defense shrimp from VP6 infection. Shrimp survival rates were significantly higher when pretreated with VA3 (∼69.0 %), Psyc (∼44.0 %), Rueg (∼43.0 %), or Halo (∼38.0 %) compared to the positive control group exposed only to VP6 (∼23.0 %). Other strains showed insufficient or adverse effects on survival. Notably, a combination of VA3, Psyc, Rueg, and Halo offered enhanced protection compared to Com12 alone, suggesting that these strains collectively provide synergistic defense mechanisms against VP6 invasion.

### Synergy effects of commensal bacteria and phage in enhancing shrimp survival against pathogenic Vibrio invasion

To evaluate whether the subset consortium comprising VA3, Psyc, Rueg, and Halo (Com4) could protect shrimp from VP6 infection and whether this protective effect could be enhanced by phage addition within the complex intestinal microbiome, we let shrimp being colonized with Com4 (5×10^6^ CFU/mL per strain) before exposing to VP6 (5×10^6^ CFU/mL) (**Fig. 5a**). then, the shrimp were maintained under standard aquaculture conditions and fed daily with phage (∼10^9^ PFU/g) throughout the experiment. Successful colonization by VP6 causes an acute infection over a five-day period with white hepatopancreas and empty digestive tracts (**Extended Data Fig. 7**), which is a typical symptom of vibriosis(*22*, *23*). Whereas shrimp with Com4 can rapidly succumb to the infection.

**Fig. 5.**
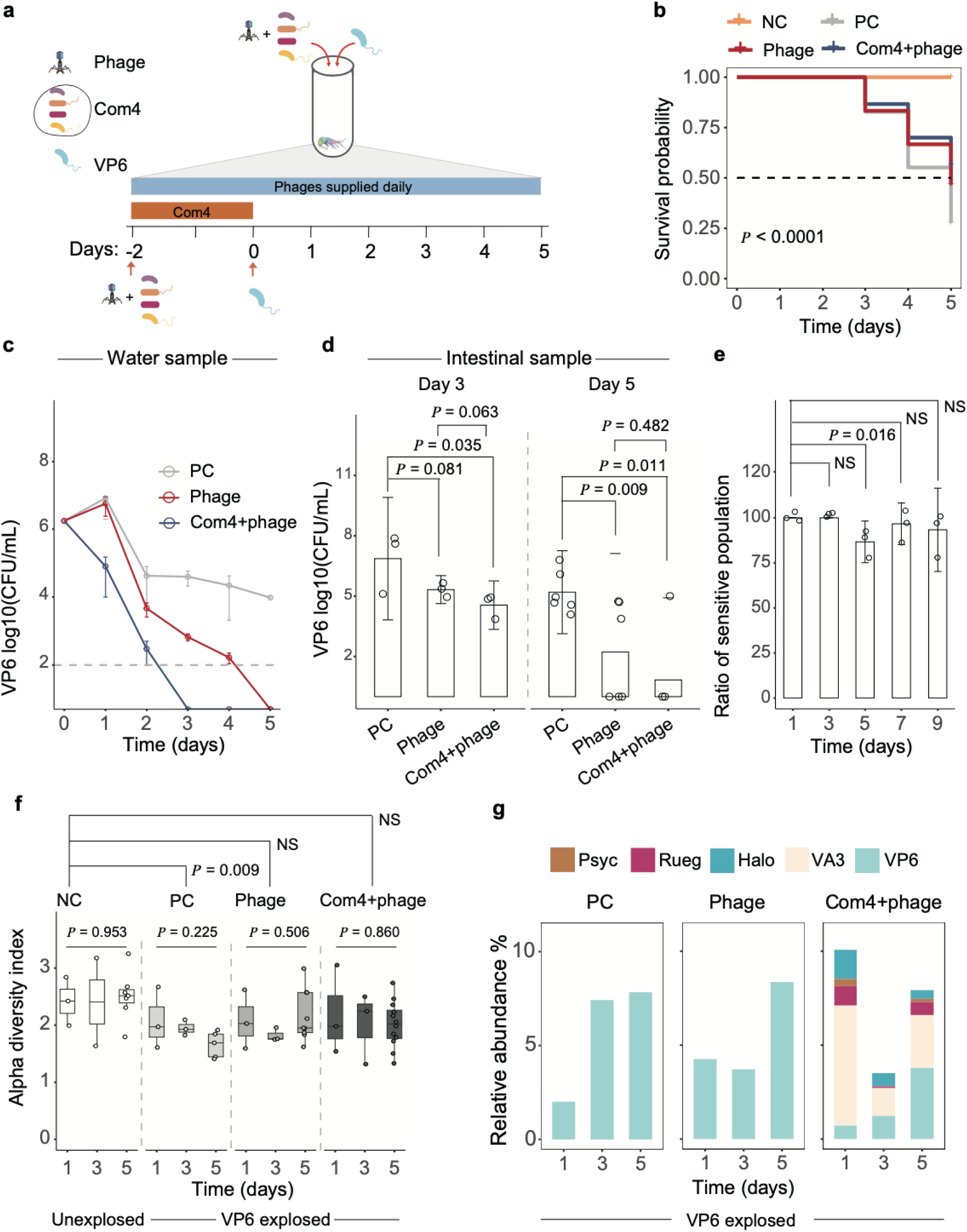
Coupled effects of phage and commensal strains on intestinal microbiota and VP6 *in vivo*. **a**, Experimental diagram for *in vivo* treatment. Shrimp was administered Com4 (5×10^6^ CFU/mL per strain) and phage for 2 days before VP6 introduction. Phage was also supplied daily in shrimp feed (∼10^9^ PFU/g) throughout the experiment. Shrimp samples were collected on days 1, 3, and 5 after VP6 exposure, and aquatic water samples were collected daily. **b**, Survival rates of shrimp over a 5-day period. Negative control: Shrimp received no treatment. Positive control: Shrimp was exposed to VP6. Phage treatment: Shrimp was treated with phage after VP6 introduction. Com4+phage: Shrimp received combined treatment with Com4 and phage following VP6 exposure. Statistical results are given as exact *P* values in brackets in the graphs. **c**,**d**, Absolute VP6 concentration aquatic water (***c***) and intestine (***d***) samples. VP6 concentrations were measured by counting colonies on selective TCBS plates. Data are presented as log10 CFU per mL for aquatic water samples and log10 CFU per g for intestinal samples. Points represent the geometric means ± SD (n = 3 ∼ 6) at different time points. Intestinal samples were collected on days 3 and 5 (n = 3 ∼ 6). In panel ***d***, a two-sided Student’s *t*-test was used. Statistical results are given in the graphs or indicated using the following: NS, not significant (*P* > 0.05, unless specified otherwise in the figure), \*\*\**P* < 0.0001. **e**, Susceptibility of VP6 isolates from shrimp intestines to the original phage. Ten VP6 isolates were collected from each shrimp sample at different time points (1, 3, 5, 7, and 9 days post-co-culture with phage). Data points represent the means ± SD of the percentage of sensitive isolates from each shrimp sample (n = 3). Significant differences between groups were assessed using pairwise Wilcox tests with adjusted *P* value. Statistical results are given in the graphs or indicated using the following: NS, not significant (*P* > 0.05, unless specified otherwise in the figure), \*\*\**P* < 0.0001. **f**, Alpha diversity of the shrimp intestinal bacterial community at day1, 3 and 5 post-infections, assessed using the Shannon index based on bacterial OTUs (> 97 % similarity). Box plots show the interquartile range with the median indicated by in line. Significant differences between groups were assessed using pairwise Wilcox tests with adjusted *P* value. Statistical results are given in the graphs or indicated using the following: NS, not significant (*P* > 0.05, unless specified otherwise in the figure), \*\*\**P* < 0.0001. **g**, The relative abundance of VP6 and the four strains of Com4 in the shrimp intestinal samples within different treatment groups. This based on 16S rRNA gene sequences and are shown.

While mortality occurred across all groups following VP6 exposure, the cumulative survival rate of shrimp in the Com4-phage treatment group significantly increased to 58% (*P* <0.001) compared to >20% in the VP6-only challenge group (Positive control) (**Fig. 5b**). The survival rate in the Com4-phage group was also notably higher than in the phage-only treatment group, confirming *in vitro* findings (**Fig. 4**) that the combination of commensal bacteria and phage more effectively inhibits VP6 invasion.

In addition, both the phage and the Com4-phage treatments effectively suppressed VP6 colonization in the aquatic environment. Plate counting assays revealed that VP6 became nearly under detectable in water surrounding the shrimp after three days in the Com4-phage treatment group, whereas similar suppression was observed only on the fifth day in the phage-only group (**Fig. 5c**). Quantitative analysis of VP6 in shrimp intestinal samples collected on day 3 and 5 showed a similar trend: VP6 abundance significantly decreased by over 90% in both treatment groups compared to the positive group (**Fig. 5d**).

Interestingly, phage and the sensitive Vibrio target (VP6) coexisted in the shrimp intestine of the phage-only treatment group. While most *Vibrio* strains isolated from intestinal samples remained susceptible to the wild type phage (**Extended Data Fig. 8**), the observed increased in phage particles alongside a decrease in Vibrio load suggested that phage-mediated suppression of VP6 was effective but limited within the intestinal environment.

### Synergistic suppression by Com4 and phage to VP6 highlighted on shrimp intestinal microbiome

To investigate the effects of the treatments on the shrimp intestinal microbiome, samples were collected at three time points for further analysis. Alpha diversity, as measured by the Shannon index, decreased in the positive control group but increased in the phage-only and Com4-phage treatment groups (**Fig. 5f**). Characterization of the microbiome revealed that both Com4 strains and VP6 successfully colonized the shrimp intestine (**Fig. 5g**). Importantly, the relative abundance of VP6 was significantly lower in the Com4-phage treatment group compared to the positive group and phage-only groups, demonstrating superior pathogen resistance and microbiome recovery in the Com4-phage treatment group.

A ternary plot of bacterial OTUs in shrimp intestinal samples demonstrated that the four commensal strains, in combination with phage predation, effectively suppressed VP6 colonization (**Extended Data Fig. 9**). These findings suggest that Com4 strains provide substantial resistance to pathogen colonization in the shrimp intestine, complementing the inhibitory effects of phage. Co-occurrence network analysis of the intestinal microbiomes in the Com4-phage group revealed positive interactions between VP6 and other *Vibrio* species, including VA3 (**Extended Data Fig. 10**) This suggests that VP6, VA3, and indigenous *Vibrio* spp. occupy similar ecological niches within the shrimp intestine, potentially contributing to complex competitive dynamics.

Together, these results highlight the synergistic effects of commensal bacteria and phage in enhancing colonization resistance against VP6. The combination of Com4 strains and phage not only improved shrimp survival rates but also restored microbiome diversity and reduced VP6 colonization more effectively than phage treatment alone. This underscores the potential of leveraging commensal-phage synergies to protect aquaculture species from pathogenic infections.

## Discussion

Pre-establishing a healthy microbiota is a pivotal strategy for enhancing resistance to pathogen colonization, primarily through mechanisms such as ecological priming(*24*) and competitive exclusion(*25*). In this study, we combined *in vitro* assays with *in vivo* shrimp models to demonstrate that phages and specific microbial species can collectively fortify microbial consortia against pathogenic invasion, thereby preventing infection. A key finding is the critical role of timing in microbial interactions, where delayed intervention relative to pathogen invasion significantly reduced the preventive effects on pathogen colonization, even with phage-mediated selective suppression. However, this limitation can be mitigated by combining phages with selected commensal species, highlighting the potential of tailored microbial consortia in enhancing pathogen resistance.

Our results further underscore the importance of microbial diversity of the gut microbiome in fortifying resistance to pathogen colonization, with more diverse microbiomes exhibiting stronger defenses against invading pathogens than less diverse communities(*3*). While many *Vibrio* species coexist harmlessly in aquatic ecosystems(*26*), certain strains, including *Vibrio parahaemolyticus* are pathogenic and cause severe disease in both animals and humans(*15*, *16*). In aquaculture, *V. parahaemolyticus* carrying the binary toxin gene pirAB is a major cause of vibriosis in shrimp, leading to hepatopancreatic damage and high mortality rates(*27*). Sequencing-based microbial profiling has demonstrated that vibriosis outbreaks correlated with alternations in the intestinal microbiome(*18*), particularly with the rapid proliferation of *Vibrio* species within the gut ecosystem(*28–30*). In our study, we observed a pronounced shift in the intestinal microbiome associated with vibriosis, characterized by a significant increase in *Vibrio* species (**Extended Data Fig. 11)**. Beta diversity analysis at the OTU level revealed clear differences between healthy and diseased shrimp, with the diseased microbiota exhibiting greater variability and lower alpha diversity, reflecting a disrupted community structure dominated by fewer bacterial taxa (pairwise Wilcoxon, *P* < 0.05) (**Extended Data Fig. 11a,b**). We identified seven key microbial biomarkers distinguishing healthy from diseased shrimp, with *Vibrio* being the predominant genus contributing to the differences in microbiome composition (**Extended Data Fig. 11c**). These findings suggest that microbial diversity may limit pathogen access to ecological niches, thereby enhancing resistance.

While microbial diversity is crucial, not all species contribute equally to colonization resistance. Keystone species, in particular, exert disproportionately strong effects, often by engaging in like competitive interactions such as nutrient competition, niche exclusion, or direct antagonism(*31*, *32*). In our study, we found that *V. alginolytic* strain VA3, a commensal strain with high nutritional overlap with *V. parahaemolyticus*, was particularly effective in outcompeting the pathogenic *V. parahaemolyticus* strain VP6. By directly competing for essential resource and ecological niches, *V. alginolytic* VA3 enhanced colonization resistance when integrated into a broader consortium, amplifying its suppressive effect on pathogen proliferation, and contributing to the overall balance of the intestinal microbiome. This highlights the potential of commensal *Vibrio* species as natural biocontrol agents, offering a promising strategy for preventing or mitigating infections in aquaculture and human health contexts.

The suppressive effects of these keystone species are further amplified when combined with phage-mediated selective suppression. In our engineered consortium, the combination of commensals with phages provided highly effective protection against the pathogenic *V. parahaemolyticus* strain VP6, which is responsible for vibriosis in our shrimp model. This synergy underscores the potential of integrating phage therapy with microbiota modulation to enhance pathogen control. Phage treatments, however, were most effective when administered early; delayed introduction of phages significantly reduced their efficacy. This attenuation may arise from coevolutionary dynamics between phages and bacteria, where phage-resistant mutants evade predation, or from phenotypic resistance in slow-growing, biofilm-forming, or stationary-phase bacterial subpopulations that escape phage attack(*33*). To mitigate this, strategies such as increasing phage density(*14*), employing phage cocktails targeting multiple receptors(*34*), or utilizing phage training approaches(*35*) have been widely considered. While these approaches show promise, they risk accelerating bacterial adaptation(*36*), potentially diminishing long-term efficacy. Alternatively, ecological control strategies, such as restricting nutrient availability(*37*) and reshaping ecological niches(*38*), represent promising approaches to complement phage therapy.

Additionally, our findings emphasize the critical role of physical separation within the shrimp intestine in limiting the efficacy of phage therapy. Consistent with previous findings(*39*) on phage-bacteria interactions in complex environments, we show that the ecological complexity, rather than phage resistance alone, is the primary factor restricting phage effectiveness. In this diverse and dynamic environment, factors such as microbial competition, resource availability, and the physical architecture of the gut create substantial challenges for phages in effectively targeting and eradicating pathogens(*33*, *40*) Given that phage-mediated bacterial killing is inherently contact-dependent(*41*), physical separation between phages and their bacterial targets, or reduced encounter rate(*42*), can significantly impact phage infection dynamics. This limitation highlights the need for integrated approaches that combine ecological control with phage therapy. For instance, combing engineered microbial consortia that occupy pathogen-preferred niches with phages tailored to specific pathogen subpopulations may yield synergistic effects, enhancing both the precision and durability of pathogen control.

Our finding also indicates that the activation of bacterial stress responses, particularly the excision of prophages, plays a key role in pathogen suppression. In response to environmental stressors, bacterial cells activate a variety of repair pathways, leading to the excision of resident prophages. The excision of prophages can disrupt the host’s competitive ability in certain environments, as the accessory genes carried by prophage often expand the host’s ecological niches(*36*, *43*). This disruption may reduce the pathogen’s growth potential, as the prophage-excised host can no longer fully exploit the resource or ecological space it once controlled. Notably, our findings underscore the importance of strain-level interactions in modulating pathogen dynamics. For example, *V. alginolyticus* VA3 was pivotal in boosting colonization resistance, both by outcompeting pathogen VP6 for critical resources and by triggering prophage activation under nutrient-limited conditions. This indicated that commensal bacteria could exacerbate nutrient limitation or alter pathogen behavior in a way that promotes prophage excision, contributing to pathogen suppression. Future experiments focusing on specific molecular pathways linking nutrient limitation, stress responses, and prophage activation would provide deeper insights into these mechanisms.

In conclusion, our study demonstrates that combining commensal bacteria with phage offers a synergistic and effective strategy for controlling pathogenic *Vibrio* infections in shrimp. This approach emphasizes the importance of understanding the interplay between phage dynamics, microbial community structure, and ecological factors in designing effective pathogen suppression strategies. By leveraging the specific suppressive effects of commensal strains and phages, this combined approach enhances disease resistance while minimizing the risk of pathogen resurgence. Taken together, our findings provide a framework for investigating interactions between phage, commensal bacteria, and pathogens, with significant potential for applications in aquaculture, as well as in human and animal health. Future research should focus on optimizing these combined interventions and expanding their applicability to other host-microbiome systems to ensure long-term stability and efficacy in pathogen control.

## Materials and Methods

### Bacterial strains and media

In this work, two media were used, 2216 marine broth (2216MB amended with 15 g/L of agar)(*44*) for commensal bacteria isolation and TCBS (Thiosulfate-Citrate-Bile-Sucrose Agar)(*45*) for vibrio selection. To ensure that each species used in this work can survive, the 2216MB was chosen as the final growth medium.

From a total collection of over 100 intestinal isolates, 12 commensal strains were selected to establish a consortium of commensal bacteria from the shrimp intestine, that could represent the healthy shrimp microbiome in subsequent experiments. The commensal bacteria collection was established by isolation from the intestine of healthy shrimp following standard isolation process(*46*). Briefly, to isolate bacteria from shrimp intestines, shrimp collected from the farm were washed gently in sterile distilled water and afterward the digestive tract was dissected aseptically. After homogenized in sterile distilled water, the suspensions were 10-fold serial diluted and placed directly on agar plates of 2216MB. In parallel, Vibrio strains were isolated from the intestine of shrimp suffering from vibriosis using selective plates TCBS. Colonies were inoculated into 3 mL of 2216MB in 15-mL culture tube overnight, shaking at 30 °C. For preservation at -80 °C, glycerol stocks were prepared by combining 500 μL of culture with 500 μL of 50 % glycerol (50 % water) in cryogenic tube. Colonies were purified by serial passage on agar plates two times. The 12 strains used in the commensal consortium were selected based on the characterization by 16S rRNA gene amplicon sequencing with the primer pair 27F (forward, 5′-AGAGTTTGATCCTGGCTCAG-3′; reverse, 5′-GGTTACCTTGTTACGACTT-3′)(*47*) representing key bacterial taxa and were whole genome sequenced (details in sequencing part). These strains belong to the genera of *Tenacibaculum*, *Algoriphagus*, *Gaetbulibacter*, *Halocynthiibacter*, *Ruegeria*, *Shewanella*, *Pseudoalteromonas*, *Psychrobacter*, *Exiguobacterium*, *Planococcus*, *Microbacterium* and *Demequina*, according to >97 % similarity of 16S rRNA genes and 500 core genes taxonomy annotation.

In parallel, *Vibrio* strains were isolated from the intestine of shrimp suffering from vibriosis using TCBS as a selective medium. The diseased shrimp were collected the same farms as above (Shanwei, Guangdong, China). From the diseased shrimp, 11 out of 100 *Vibrio* isolates with relatively distant phylogenetic relationships were picked as candidates for further study based on 16S rRNA gene also and two representative strains were further whole genome sequenced.

### Whole genome sequencing of bacterial strains

For whole genome sequencing, overnight culture of each strain was concentrated by centrifugation at 8,000 rpm, 10 min. DNA extraction, purification, and density estimation were made by the protocol of bacteria DNA extraction kit without modification (referred to previous description). These DNA samples were sequenced by the Illumina Nova-PE250 sequencer platform (Annoroad gene technology Co. Ltd, China). The high-quality filtered reads were assembled using SOAP de novo v2.40(*48*) by default parameters. Complete genome sequences were manually checked.

### Virulence gene determination

The *Pir* virulence gene has been known to encode PirAB toxins that cause acute hepatopancreatic necrosis disease (AHPND)(*18*, *49*). To determine the pathogenicity for these 11 *Vibrio* strains, DNA of each *Vibrio* strains was extracted and used for *pir* gene detection with the specific primers (forward, 5′-ATGAGTAACAATATAAAACATGAAAC-3′; reverse, 5′-GTGGTAATAGATTGTACAGAA-3′)(*50*). The PCR products were detected using agarose gel electrophoresis with 250bp DNA ladder (DL10K Plus DNA Marker).

### Phage isolation and characterization

Using *Vibrio* VP6 as host bacteria, phages were isolated from the sewage of shrimp farms (Shenzhen, China) using the plaque assay as described previously(*20*). For the phage sequencing, concentrated phage particles were treated using DNase I and RNase A (New England BioLab, England) to remove bacterial nucleic acids. Then, the phage genomic DNA was extracted using a lambda bacteriophage genomic DNA Rapid extraction kit (DN22; Aidlab, China). To determine the host-ranges of the specific phages against all other strains used in this work, we tested the bacterial susceptibility by a spot titer technique. Bacterial strains were grown in 5 mL of 2216MB overnight from single colonies streaked on 1.5 % agarose plates supplemented with 2216MB. Phage lysates were prepared in advance. Ten µL drop spots of phage lysates with ∼10^9^ PFU/mL (or phage buffer as a control) were pipetted onto the bacterial lawn made using double-layer method and incubated at 30 °C in air incubator overnight for phage plaque. For some species, the plates were kept for 48 h. The phage (OM319461) with a high specificity to VP6 was selected in this study.

### Consortium preparation and invasion *in vitro*

Each commensal strain selected for the consortium was cultured independently in 2216MB. To assemble the consortium, the culture of each strain was diluted to OD600 = 0.01 and then all strains were mixed equally in a final volume of 200 μL. This consortium of 12 strains (Com12) was then cultured using 100-well plates at 30 °C with shaking (Bioscreen C, medium amplitude) for 48 h. Subsamples for the quantification of colony forming units (CFU) of the consortium components were collected at 5 timepoints, including 6h, 12h, 24h, 36h and 48h. 16S RNA amplicon sequencing was adopted to measure the dynamics. CFU of individual strains was determined using a culture-independent method(*51*), in which an interior label was utilized. For the interior label, a specific number of *Escherichia coli* strain MG1655 cells (1.21×10^7^ cells, equivalent to 3.03×10^8^ CFU/ml) was added into each sample immediately before DNA extraction. In parallel to control growth experiment with Com12, incubations with addition of *Vibrio* isolates were established to simulate a *Vibrio* pathogen invasion in the assembled consortium and carried out under the same condition.

An experiment to simulate the *Vibrio* pathogen invasion and afterward with phage treatment in the assembled consortium was carried out accordingly. In detail, phage lysate (∼10^11^ PFU/mL) with a specific MOI = 1 was added into the consortium at the five specific serial time points, including 0 h, 2 h, 6 h, 12 h and 24 h. To simulate the prophylactic effect, phage lysate was added into the consortium (Com12) initially and then the pathogenic *Vibrio* strain was added at six serial time points, including 2 h, 6 h, 12 h, 24 h and 36 h. In parallel, another group without phage addition was set up as control. Furthermore, to investigate the effect of extra addition of commensal *Vibrio* VA3 based on the phage treatment with the timing as the variables, the specific culture of VA3 was introduced into the assembled consortium, with VP6 invasion and phage addition performed accordingly. A group without phage addition was set up as control. All experiments were carried out under the same condition. At each time point, samples were collected and stored at -80 °C immediately for DNA extraction and the 16S rRNA gene V4 region amplicon sequencing.

## Animal procedures and sampling

### Comparison of intestinal microbiomes between diseased and healthy shrimp

Pacific white shrimp, *Penaeus vannamei*, were collected from the aquafarm (Shenzhen Alpha feed Agriculture and Animal Husbandry Co., Ltd; Shan-Wei, China). Six healthy shrimp and six diseased shrimp with symptoms of *Vibrio* infection were recruited to compare the intestinal microbiome. The healthy shrimp showed a normal size hepatopancreas with dark orange color and a full stomach and midgut. The diseased shrimp were shown with pale, atrophied hepatopancreas and an empty digestive tract. Shrimp bodies were sterilized with 75 % alcohol and the intestine dissected out in an ice-cold and sterile environment. The midgut was washed 5 times in sterile PBS and put into a sterile tube containing 1 ml sterile PBS buffer and 3 mm sterile grinding steel ball. The mixture was homogenized thoroughly by a tissue crusher (60 Hz, 1 min). DNA extraction was immediately implemented for these homogenized samples.

### The feeding and administration procedure for experimental shrimp

The healthy shrimp larvae were obtained from Hongzheng Aquatic Product Limited Company (Zhanjiang, China) with 21 days of breeding after hatching. Shrimp were routinely fed and maintained in a lab environment. In brief, shrimp larvae were fed twice per day and maintained in artificial seawater (12 ‰ salinity, 28 °C). These shrimp without endogenous *Vibrio* pathogens were verified by PCR and culturing on TCBS.

a. *Vibrio* pathogenicity evaluation. To assess the pathogenicity of *Vibrio* strains VP6 and VA3, the strains were cultured from single colonies at 30 °C overnight and concentrated by centrifugation at 8,500 rpm for 10 min. The concentration of the re-suspended bacteria was quantified using TCBS plates. Healthy shrimp were randomly divided into two groups and the bacterial solution was added and mixed for a final concentration of 5×10^6^ CFU/mL. These shrimp were maintained at 28 °C with continuous oxygenation. The pathogenicity of these strains was assessed according to the post-infection mortality of the shrimp for 5 days.
b. Protection of the commensal consortium and monospecies on shrimp. To avoid the potential effect of the intrinsic microbiome, shrimp were subjected to Florfenicol (50 mg/L, aladdin#E2018032), Ciprofloxacin (50mg/L, MACKLIN#C11407180), and Sulfamethazine (25 mg/L, aladdin#C2003138) for 48 h at 28°C. Subsequently, sterile artificial seawater was used to wash away the residual antibiotics three times. The sterile shrimp were then exposed to either the defined consortia (Com12, -/+VA3) or by each commensal strain individually, at a final concentration of 5×10^6^ CFU/mL. In brief, frozen stocks of individual strains were streaked out on marine broth, single colonies were picked, and each strain was cultured separately in 15 ml Falcon tubes for 24 h (some for 48 h). Cultures were pelleted and washed twice and re-suspended in sterile artificial seawater. For the consortium, different strains were then mixed equally. After bacteria recovery for 48 h, pathogenic *Vibrio* was added with a final concentration of 5×10^6^ CFU/mL. For comparison, two control groups were set up: positive control where shrimp were exposed to the pathogenic *Vibrio* without the establishment of commensal strains; negative control, without the addition of the pathogenic *Vibrio* strain. In addition, to quantifying the preventive effects of the consortium, we investigated the protective effect of each of the individual Com12 strains. In that experiment, each commensal bacterial strain was administered to randomly divided shrimp groups separately at the same condition. After bacteria exposure for 48 h, pathogenic *Vibrio* was added at a final concentration of 5×10^6^ CFU/mL, positive (pathogen but no commensal) and negative (commensal but no pathogen) control groups as described above.
c. Efficacy Assessment *in vivo*. Shrimp were used to test the combined suppressive efficacy of the phage and commensal bacterial strains on the pathogenic *Vibrio* strain in shrimp. First, we ensured the absence of pathogenic *Vibrio* strain (VP6) in the shrimps’ intestine, using the PCR method to screen for the *pir* gene and the *Vibrio* selective TCBS plates to culture the intestinal microbes, we were unable to identify them. Shrimp were randomly and equally divided into four groups. For two of these groups, shrimp were pre-exposed to phage and phage-commensal combination with final concentrations of 5×10^7^ PFU/mL and 5×10^6^ CFU/mL, respectively; after two days, the pathogenic *Vibrio* strain was inoculated into these two groups with a final concentration of 5×10^6^ CFU/mL. In addition, phage was provided to the shrimp by mixing with feed (10^9^ PFU per gram) daily. The other two groups were established as a positive control that was exposed to the pathogenic strain and negative control that was amended with sterile PBS instead, respectively. Shrimp were sampled on the 1st, 3rd and 5th days, respectively.
d. Quantitative determination of the pathogen and the phage *in vivo*. Shrimp were inoculated with VP6 and the phage simultaneously. Shrimp were sampled on the 1st, 3rd, 5th, 7th and 9th days, respectively. The samples were homogenized and the concentration of VP6 was determined by counting the number of VP6 colonies on the TCBS plate. For the phage concentration, the sample homogenate was centrifuged at 8,500 rpm for 10 min and filtered by a 0.22 μm filter. Then the filtrate was serial diluted and 10 μL diluent was added to marine broth agar containing 200 μL culture of VP6. In addition, 30 green colonies were picked from the TCBS agar plate into 1ml marine broth per sampling time-point to test the bacterial sensitivity to the original phage. In brief, 10 μL of corresponding cultures and 10μL phage lysis (∼10^11^ PFU/mL) were inoculated into 180 μL marine broth in a 100-well plate. The plate was incubated at 30 °C with shaking (medium amplitude) for 12 h. Phage sensitivity of the isolated VP6 strains was determined by the growth curve.

PBS buffer washing was used to disinfect the shrimp surface, and intestinal samples were aseptically dissected as described previously(*19*). For DNA extraction, the intestinal sample was put into a sterile tube containing 1 mL sterile PBS buffer and 3mm sterile grinding steel ball and was homogenized thoroughly by tissue crusher (60 Hz, 1 min). DNA extraction was immediately implemented for these homogenized samples. The density of the pathogenic *Vibrio* VP6 was determined by the presence of the specific colony, which had green color on TCBS plates after 12h incubation at 30 °C. Three technical replicates were used for each measurement.

### Spent media assay

To quantify the pairwise positive and negative effects between 11 selected individual commensal bacteria and 2 presentive *Vibrio* strains, each bacterial isolate was grown in conditioned medium from each of the other bacteria for quantification of growth promotion or inhibition. Conditioned Medium Preparation was conducted using the method by Marjoin *et al*(*21*). The spent medium was prepared by culturing isolates in a marine broth medium, followed by centrifugation and filtering. To prepare the conditioned medium, the marine broth spent medium fraction was mixed with the marine broth medium. The spent medium fraction was replenished with marine broth constituents, so that the concentration in the final conditioned medium ranged from 0.6× (for constituents that were consumed in the spent medium) to 1.0× (for constituents that were not consumed in the spent medium) of the concentration in marine broth. The pH of the conditioned medium was adjusted to 7.6, which is consistent with the fresh marine broth medium. Each isolate was incubated for 36 h in one well of a 96-well plate containing 200 μL of pre-warmed conditioned medium or complete medium. Bacteria cultures were inoculated by ∼0.2 μL from a thawed overnight culture stored at −80 °C. Plates were incubated at 30°C and shaken at 200 rpm in aerobic conditions. OD600 was recorded every ∼30 min. Quantification of the effects was performed based on the ratio of the maximum OD600 in conditional medium comparing to in the fresh medium according to the method by Marjoin *et al*(*21*).

### Prophage quantification

The relative production of prophage (Vpp2) in conditioned medium or co-culture were quantified using qPCR as studied previously(*52*) with some modifications. Briefly, the supernatants of *Vibrio* VP6 cultured in different conditions were collected, treated with DNase to remove residual DNA and incubated for 30 min at 95 °C to open phage capsids for releasing phage DNA. Digestive solution was 10-fold diluted as template for quantitative PCR. Bacteria and prophage specific primers were used, and the qPCR procedure were following the protocol of Luna Universal qPCR kit (NEB). The relative expression quantity of prophage Vpp2 was analyzed by comparing the relative amplification within samples of phage-specific primer pairs to host-specific primer pairs *gyrB* (**Extended Data Table 3**).

### Competition assay

To investigate the competitive inhibition of between VA3 and VP6, we cocultured these two strains together in 96-well plate and quantified their growth by plate reader. In detail, overnight cultures of VA3 and VP6 were adjusted to the same OD600 by fresh medium and subsequently transferred into a new culture equivalently. The OD600 value was measured by the plate reader in shaking mode. This assay was performed with three biological replicates. CFU concentration of VA3 and VP6 was quantified using standard plate counting method with 10-fold gradient dilutions, because the indication that VA3 and VP6 displayed distinct colony characteristics to each other on morphology on selective agar plate, such as size and color.

### High throughput sequencing of 16S rRNA gene

Total DNA of shrimp intestine and bacterial community in vitro was respectively extracted using TIANamp Marine Animals DNA Kit (TIANGEN, China) and TIANamp Bacteria DNA Kit (TIANGEN, China) following the manufacturer’s directions. The concentration and purity of total DNA were determined by Nanodrop. The primer pair of 515F (5’-GTGCCAGCMGCCGCGGTAA-3’) and 806R (5’-GGACTACHVGGGTW TCTAAT-3’)(*53*) was used to amplify the V4 region of 16S rRNA gene, which was modified with a barcode tag containing a random 6-base oligo. Sequencing libraries were generated using DNA PCR-Free Sample Preparation Kit (Illumina, USA) and the library quantity was determined by Qubit 2.0 Fluorometer (Thermo Scientific, USA). Libraries were sent for sequencing by a Hiseq2500 platform (Illumina, USA), which was conducted by Novogene Bioinformatics Technology Co., Ltd. (Beijing, China). Raw data generated from Hiseq2500 platform were paired-end reads.

### Data analysis

Tables of taxonomic relative abundance were generated from Fastq files using the standard DADA2 v1.34.0(*54*) pipeline using paired-end reads. Per the pipeline, quality score plots were inspected to determine inflection points for drop-offs in the quality of forward and reverse reads and reads were truncated at these points; these points were position 150 and 200 for forward and reverse reads, respectively. Sequences were assigned to OTUs (Operational Taxonomic Units) with 97 % similarity. For time-series samples, all retained reads from the pipeline after standard filtering unambiguously corresponded to the strains in the defined consortium. As for shrimp intestine samples, the refined reads were aligned using SILVA SSU Ref NR database release 132, December 2017(*55*). PhyML v3.3(*56*) was then used with default parameters to build a maximum likelihood phylogeny from the alignment. The median read depth of shrimp intestine samples data was 12362 ± 5436 reads and for time-series data was 35932 ± 24616 reads. The 16S rRNA gene analysis was performed using QIIME2 (https://docs.qiime2.org/2019.1/)(57, *58*). Bray-Curtis distance was used to evaluate the strain complexity differences of samples.

Co-occurrence network of Com4 and VP6 correlated with others in the shrimp intestinal microbiome was constructed based on OTU abundance data using method as described previously(ref). Pairwise correlations were computed using Pearson’s correlation coefficient to assess the relationships between OTUs. An adjacency matrix was then constructed, with edges representing significant correlations. The resulting network was visualized using Cytoscape v3.10.2(*59*).

Bray-Curtis dissimilarity was calculated using the relative abundance of each bacterial strain in longitudinal phage perturbation experiments using the vegan package in R. The Bray-Curtis dissimilarity was defined between different shrimp at the same time point, and between time points for different treatments. Cluster Analysis and Principal Coordinates Analysis were used to generate graphical differences in community composition. Boruta(*60*) and Random Forest(*61*) for feature selection were used to identify biomarkers between shrimp groups.

One-way ANOVA with the Kruskal-Wallis test was performed. MRPP, ANOSIM, and PERMANOVA were conducted to statistically test whether there is a significant difference between two groups with vegan package (using the function adonis) in R. Permutational analysis of multivariate dispersions (PERMDISP) was for the analysis of multivariate homogeneity of group dispersions by Vegan package in R (https://cran.r-project.org/web/packages/vegan/).

Genomic analysis was applied to group proteins into pathway families. To calculate protein overlap among the microbial community, we performed a genomic-based analysis to identify shared protein families across the species within the community. First, genome sequences of all species in the community were obtained from sequencing efforts in this study. These genomes were annotated using the RAST server(*62*) to assign functional annotations and predict protein-encoding sequences. Proteins were then clustered into orthologous groups using the OrthoMCL v2.0.9(*63*) algorithm based on sequence similarity with the default parameters. Protein overlap was calculated by determining the intersection of orthologous groups across all genomes in the community. For each protein family, the presence or absence across species was recorded, and a protein matrix was generated. The overlap index for each species was defined as the proportion of shared protein families relative to the total number of protein families in the community.

All statistical analyses were conducted in the R open-source software (R core team 2025). All statistical tests were two-sided, and differences were considered statistically significant if the *P* value < 0.05. In particular, the stats packages were used to run all ANOVAs and Mann-Whitney U test (using the *lm* and Wilcox test functions), the vegan package was used to calculate the Alpha diversity (using the diversity function, Shannon index) and Patterns of similarity among samples (using the metaMDS function). Feature selection to identify the driver strains was performed using Boruta in the Random Forest package and the same name package with default parameters. The Boruta algorithm was used to select the taxa with significant predictive power as ranged by the importance in packages Boruta. To visualize the difference between different treatment groups, the ggtern package was used for the ternary plot. Statistical details of experiments can be found in the Results section and the figure legends.

## Data availability

The genome sequences of all bacterial strains in this study have been deposited at the BioProject No: PRJNA854744, under accession numbers SAMN29790832 (*Vibrio* VP6), SAMN29790135 (*Vibrio* VA3), SAMN29790134 (*Tenacibaculum* sp.), SAMN29790133 (*Shewanella* sp.), SAMN29790127 (*Ruegeria* sp.), SAMN29790108 (*Pseudoalteromonas* sp.), SAMN29790107 (*Planococcus* sp.), SAMN29790105 (*Microbacterium* sp.), SAMN29790062 (*Halocynthiibacter* sp.), SAMN29639023 (*Demequina* sp.), SAMN29639016 (*Algoriphagus* sp.), SAMN29637731 (*Exiguobacterium* sp.), SAMN29446561 (*Psychrobacter* sp.), and SAMN30247027 (*Gaetbulibacter* sp.). The accession number for raw 16S rRNA gene sequencing data reported in this study is deposited in the Sequence Read Archive (SRA) of the National Center for Biotechnology Information (NCBI): Healthy shrimp 1∼6 (SRR22750841, SRR22750923, SRR22750839, SRR22750840, SRR22750838, SRR22750842), Diseased shrimp 1∼6 (SRR22750925, SRR22751313, SRR22751216, SRR22750929, SRR22750924, SRR22750927). In vitro experiment 1∼5 (SRR22750926, SRR22751312, SRR22751214, SRR22751215, SRR22750928). The sequence accession numbers of phage used in this study is OM319461.

## Code availability

The present study did not generate code, and mentioned tools used for the data analysis were applied with default parameters unless specified otherwise.

## Acknowledgments

This work received support from the Strategic Priority Research Program of the Chinese Academy of Sciences (No. XDB29050500); Shenzhen Institute of Synthetic Biology Scientific Research Program (Grant no. JCHZ20200001). MM was financially supported by the European Union under the Horizon Europe Programme, Grant Agreement No. 101084204 (Cure4Aqua), and by the Innovation Fund Denmark (AQUAPHAGE project).

## Contributions

Y.M., L.C. conceived the project. Y.M., L.C. and Z.H. designed the pipeline. Z.H. performed the *in vitro* assays with the assistance from L.C, and conducted *in vivo* shrimp trails. L.C. designed and carried out bioinformatic analyses with inputs from Z.H., and performed the statistical analyses. C.L. and M.M. provided constructive suggestions. L.C. prepared the figures with inputs from Z.H., and wrote the first draft of the manuscript together with Y.M. and Z.H., and input from all the authors. This manuscript was then edited by all the authors. L.C., Y.M. and Z.H. supervised the generation of the manuscript.

## Corresponding authors

Correspondence to Yingfei Ma.

## Extended Data

**Extended Data Fig. 1** One-step PCR detection of the PirAB toxin gene in 11 *vibrio* strains using the specific pair primers AP3-F and AP3-R(*50*). The PCR targeted gene is *pirA*^VP^, with amplicon size is 333bp. Marker with 250bp DNA ladder was used to evaluate the fragment size.

**Extended Data Fig. 2** Morphological characterization of the shrimp (Penaeus vannamei) exposed to Vibrio strains VA3 and VP6. Control group: shrimp without treatment; VA3 group: shrimp with VA3 treatment; VP6 group: shrimp with VP6 treatment.

**Extended Data Fig. 3** Stacked bar plot of the microbial dynamics within the consortium (Com12). The consortium consisting of 12 isolated commensal bacterial strains was cocultured with different *Vibrio* strains, respectively. Sampling was conducted at 6 h, 12 h, 24 h, 36 h, and 48 h. The text below each subplot denotes the *vibrio* strain used to invade the consortium. Control: the consortium without vibrio invasion. The absolute concentration of each strain was assessed using a marker (*E.coli* MG1655) with a specific concentration for sequencing. More detail can be checked in the methods.

**Extended Data Fig. 4** Transmission electron micrograph (TEM) and genomic annotation of phage VP6phageC. TEM reveals a myoviral morphology. Scale bar (in white), 50 nm. **d**, Annotated genome of phage VP6phageC colored based on detected molecular functions of individual gene products. The arrow represents the gene length and its transcript direction.

**Extended Data Fig. 5** Growth curves of VA3 and VP6 based on optical densities (OD600 value). The OD600 value was measured using an automated 96-well plate reader in shaking mode. This assay was performed with three biological replicates of each stain and only the mean values and error bar were displayed.

**Extended Data Fig. 6** Relative quantities of prophage excision when cultured within different conditions. This ratio was evaluated using qPCR quantification the number of prophage-specific gene, to the single-copy chromosomal gene *gyrB*. Unless otherwise noted, experiments were performed with three biological replications. Significant difference was evaluated by Wilcox test. \*\*\**P* < 0.001 compared to control.

**Extended Data Fig. 7** Morphological characterization of the shrimp (*Penaeus vannamei*). Negative control: shrimp without treatment showed a black-brown hepatopancreas, full stomach and midgut. Positive control: shrimp with VP6 treatment showed a light color hepatopancreas, empty stomach and mid-gut.

**Extended Data Fig. 8.** Dynamic changes of VP6 (CFU/g intestinal sample) and the phage (PFU/g intestinal sample) in the shrimp intestine. Data points represent the means ± SD of VP6 and the phage in the shrimp samples (n = 3) collected at different time points. Absolute concentrations in the y-axis are shown in the log10 scale.

**Extended Data Fig. 9** Ternary plot of 194 taxa in shrimp intestine to evaluate and decipher the difference effect between different treatments. Each variant represents microbial species (OTU level) and colored by phylum (except VP6). The size was proportional to the average abundance for each taxon.

**Extended Data Fig. 10** Co-occurrence network of Com4 and VP6 correlated with others in the shrimp intestinal microbiome. Blue and gray edges indicate significant negative and positive correlations, respectively. To highlight the important correlations, the significant level was set up with *P* value < 0.001 and R-square threshold 0.5. Nodes represent microbial OTUs (species level) and colored on the phylum level (except for the Com4 strains), with sizes proportional to the number of edges connected to the OTUs. The edge information was shown in Table S3.

**Extended Data Fig. 11** Characterization of the intestinal microbiome of shrimp **a**, Between-group principal coordinate analysis (PCoA, axes 1 and 2) for the intestinal microbial composition data (16S rRNA) of shrimp. PCoA analysis was based on Bray-Curtis dissimilarity with default parameters. Sample points, from the diseased (n = 6, in pink) and healthy (n=6, in blue) shrimp, are marked by colors according to the groups and annotated via ellipses with 95% confidence. The significant difference (*P*-values) between groups was analyzed by the Adonis function using the distance matrices and the permutation test with pseudo-*F* ratio. **b**, Alpha diversity by the Shannon index was compared using the Wilcox-test. The significant difference is donated by asterisks (*, *P* value< 0.05). **c**, Discrimination of biomarkers among the top 92 genera using Random Forests with Boruta feature selection. Different colors and directions display the relative abundance of these biomarkers in diseased (pink colored) and control healthy (blue colored) shrimp. The circle scale represents the importance value (from 1.5 to 5.1) contributing to the discrimination between two groups.

**Extended Data Table 1** Taxonomy information of 14 species of the synthetic intestinal microbiome consortia.

**Extended Data Table 2** Spot tests of phages against each of the isolated commensal bacterial strains in this study. 10 μL of high dense of phage lysate (∼10^10^ PFU/mL) of VP6phageC was spotted onto lawns of each tested strains and incubated for 24 h or 48 h. Black/gray filling indicates whether clear zone observed or not on the bacterial lawn.

**Extended Data Table 3** Primer list used for qPCR assay. The prophage-specific primers (pro_VPP2) amplify a region associated with phage replication, while the internal control primers target the *gyrB* gene to normalize expression levels.

