## supplementary figures and tables for "Synergistic effects of commensals and phage predation in combating pathogen infections: a shrimp model"

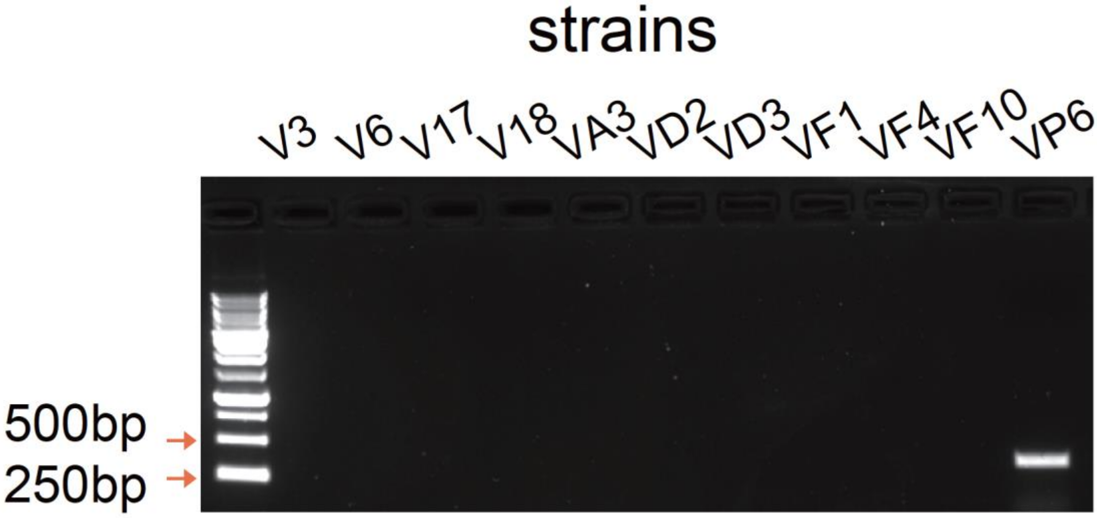


**Extended Data Fig. 1.** One-step PCR detection of the PirAB toxin gene in 11 *vibrio* strains using the specific pair primers AP3-F and AP3-R(*1*). The PCR targeted gene is *PirA*$^{VP}$, with amplicon size is 333bp. Marker with 250bp DNA ladder was used to evaluate the fragment size.

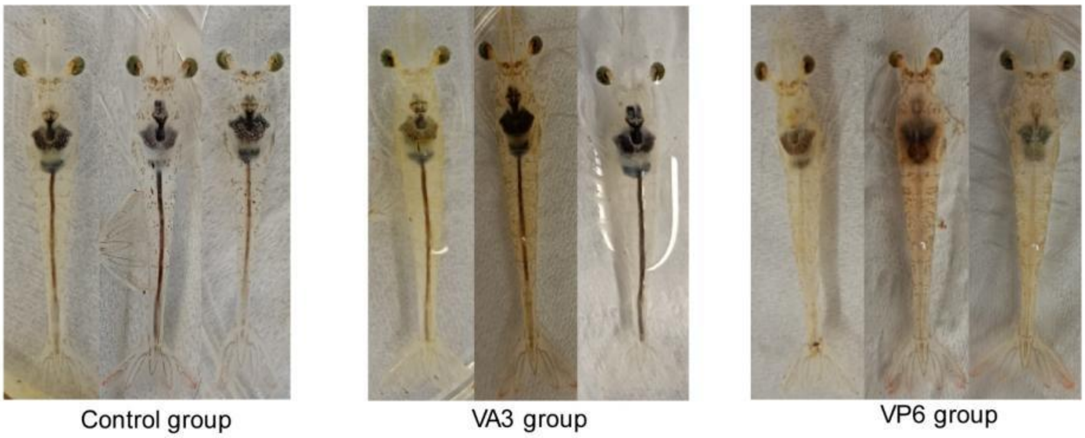


**Extended Data Fig. 2.** Morphological characterization of the shrimp (Penaeus vannamei) exposed to Vibrio strains VA3 and VP6. Control group: shrimp without treatment; VA3 group: shrimp with VA3 treatment; VP6 group: shrimp with VP6 treatment.

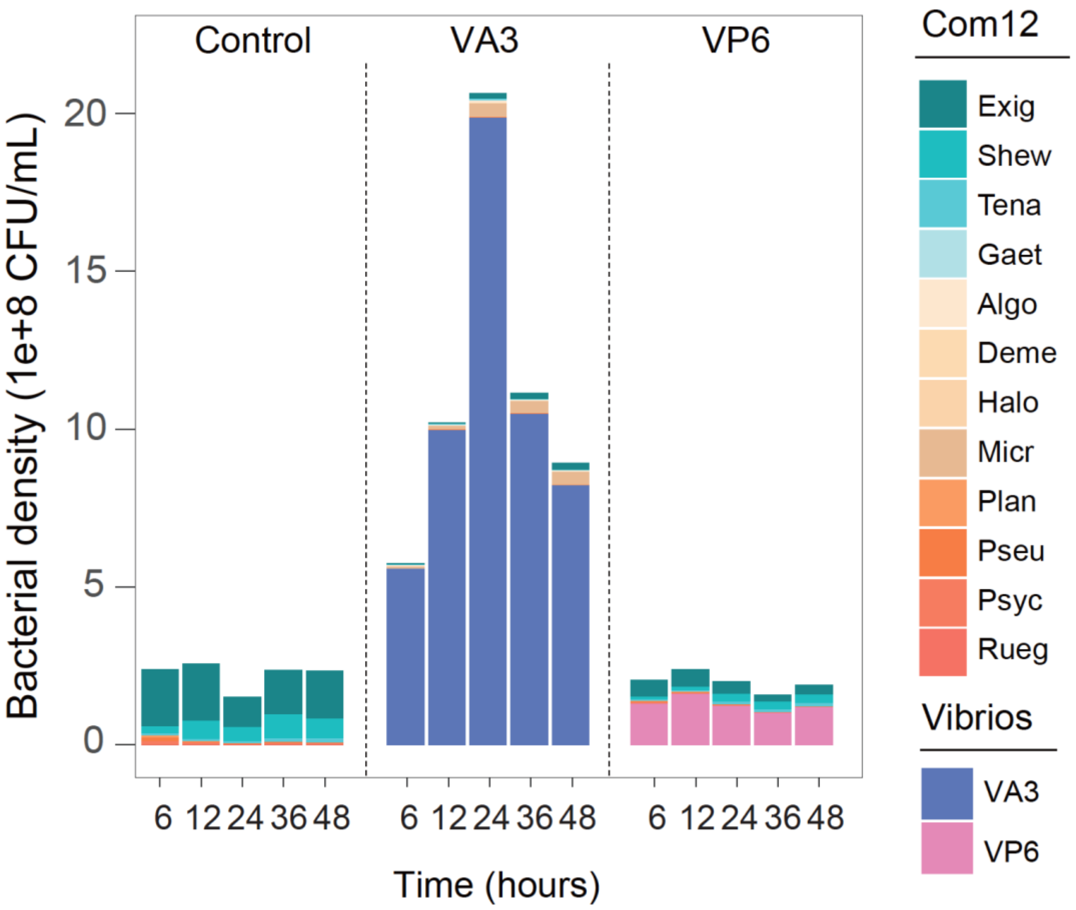


**Extended Data Fig. 3.** Stacked bar plot of the microbial dynamics within the consortium (Com12). The consortium consisting of 12 isolated commensal bacterial strains was cocultured with different *Vibrio* strains, respectively. Sampling was conducted at 6 h, 12 h, 24 h ,36 h, and 48 h. The text below each subplot denotes the *vibrio* strain used to invade the consortium. Control: the consortium without vibrio invasion. The absolute concentration of each strain was assessed using a marker (*E.coli* MG1655) with a specific concentration for sequencing. More detail can be checked in the methods.

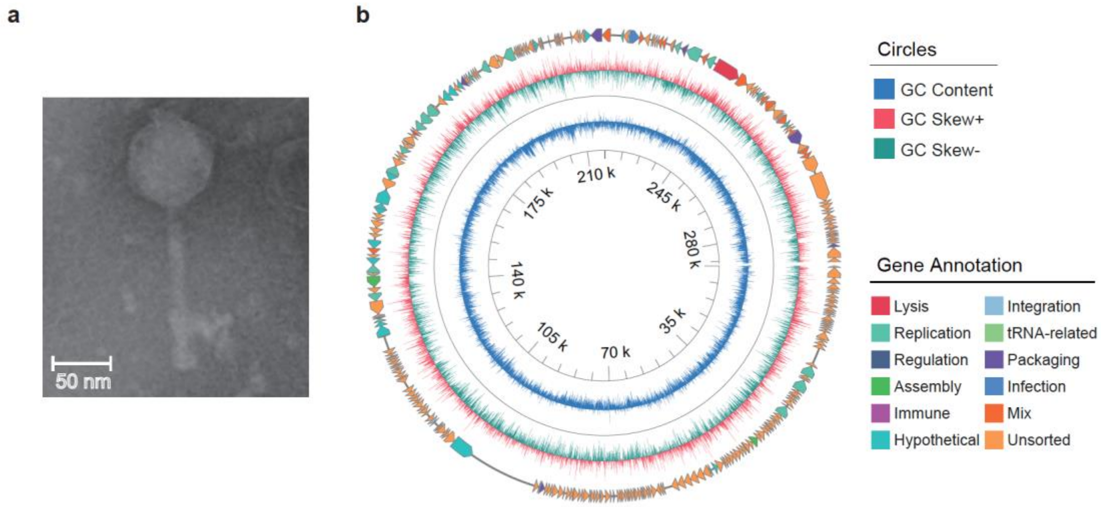


**Extended Data Fig. 4.** Transmission electron micrograph (TEM) and genomic annotation of phage VP6phageC. TEM reveals a myoviral morphology. Scale bar (in white), 50 nm. **d**, Annotated genome of phage VP6phageC colored based on detected molecular functions of individual gene products. The arrow represents the gene length and its transcript direction. This genome was visualized GenoVi(*2*).

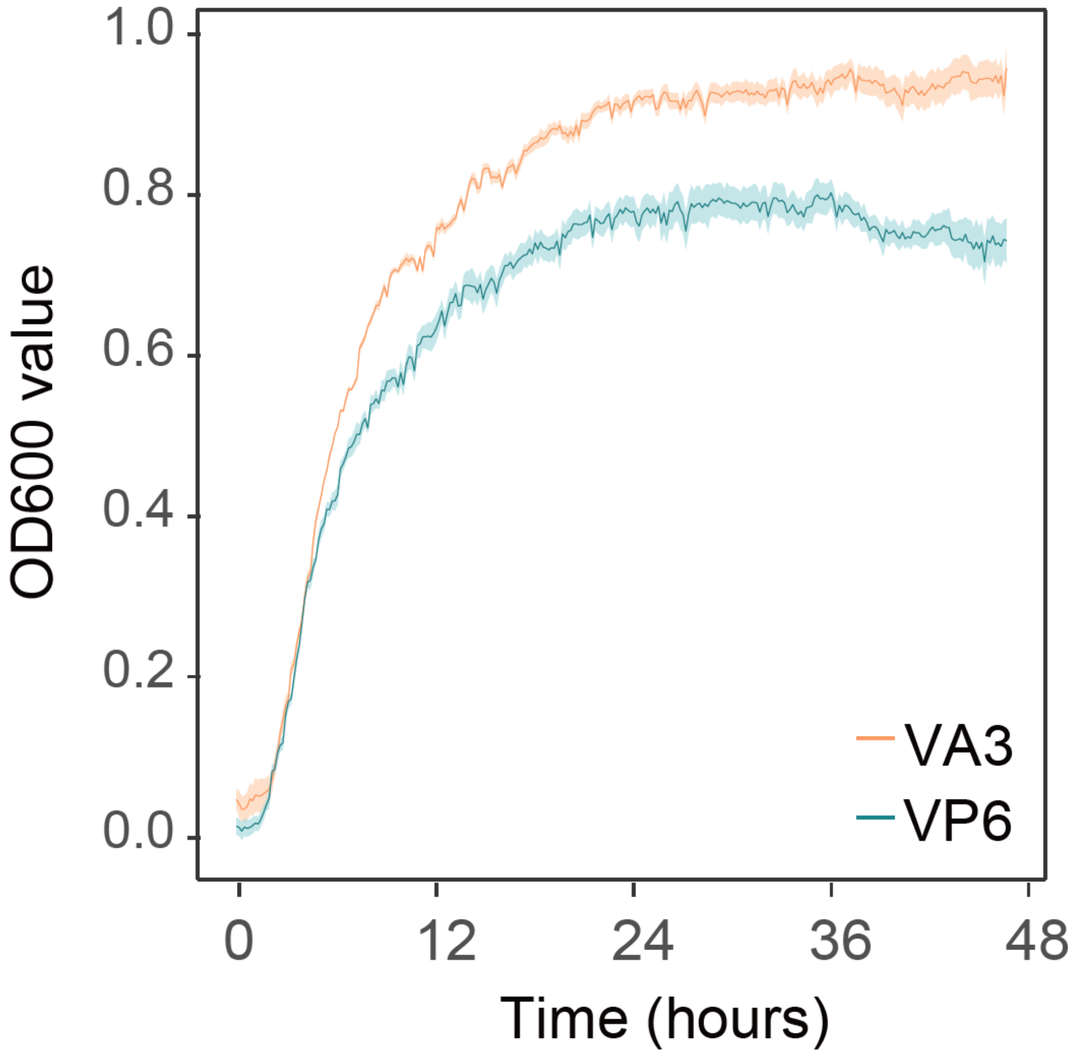


**Extended Data Fig. 5.** Growth curves of VA3 and VP6 based on optical densities (OD600 value). The OD600 value was measured using an automated 96-well plate reader in shaking mode. This assay was performed with three biological replicates of each stain and only the mean values and error bar were displayed.

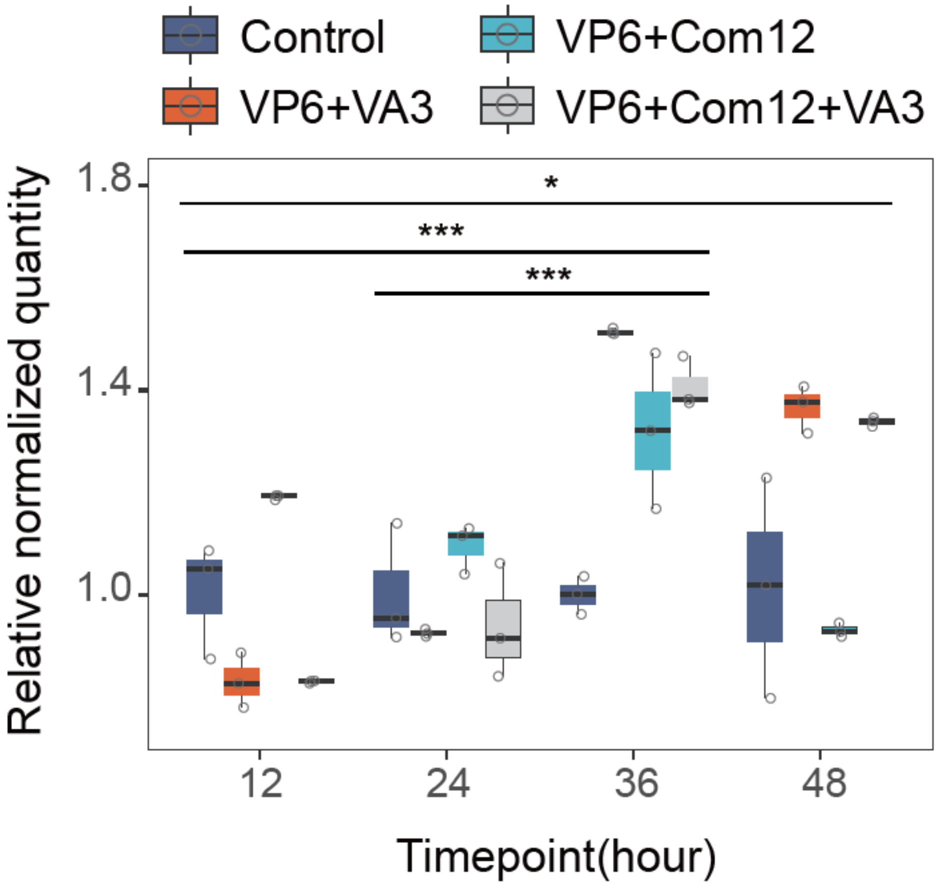


**Extended Data Fig. 6.** Relative quantities of prophage excision when cultured within different conditions. This ratio was evaluated using qPCR quantification the number of prophage-specific gene, to the single-copy chromosomal gene *gyrB*. Unless otherwise noted, experiments were performed with three biological replications. Significant difference was evaluated by Wilcox test. ***$P < 0.001$ compared to control.

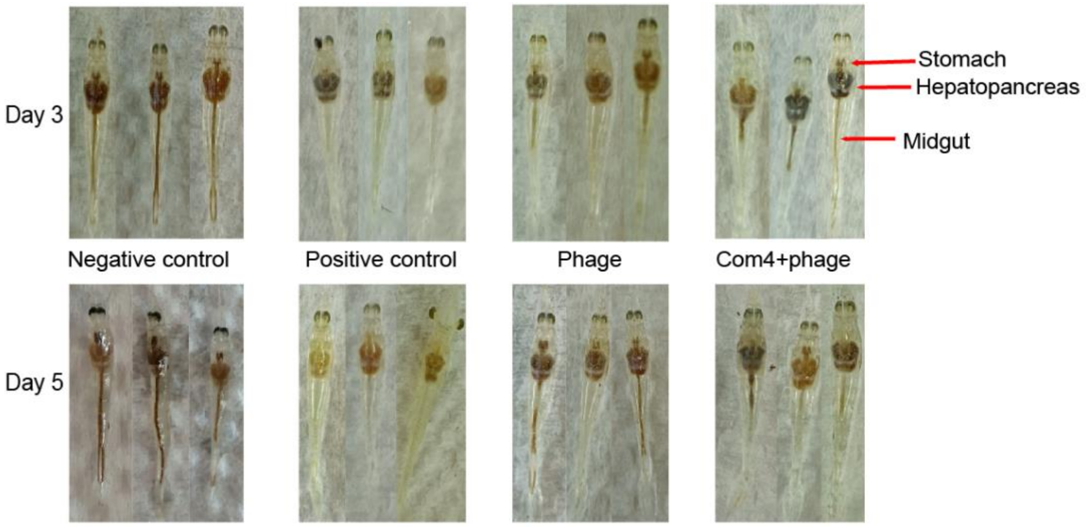


**Extended Data Fig. 7.** Morphological characterization of the shrimp (*Penaeus vannamei*). Negative control: shrimp without treatment showed a black-brown hepatopancreas, full stomach and midgut. Positive control: shrimp with VP6 treatment showed a light color hepatopancreas, empty stomach and mid-gut.

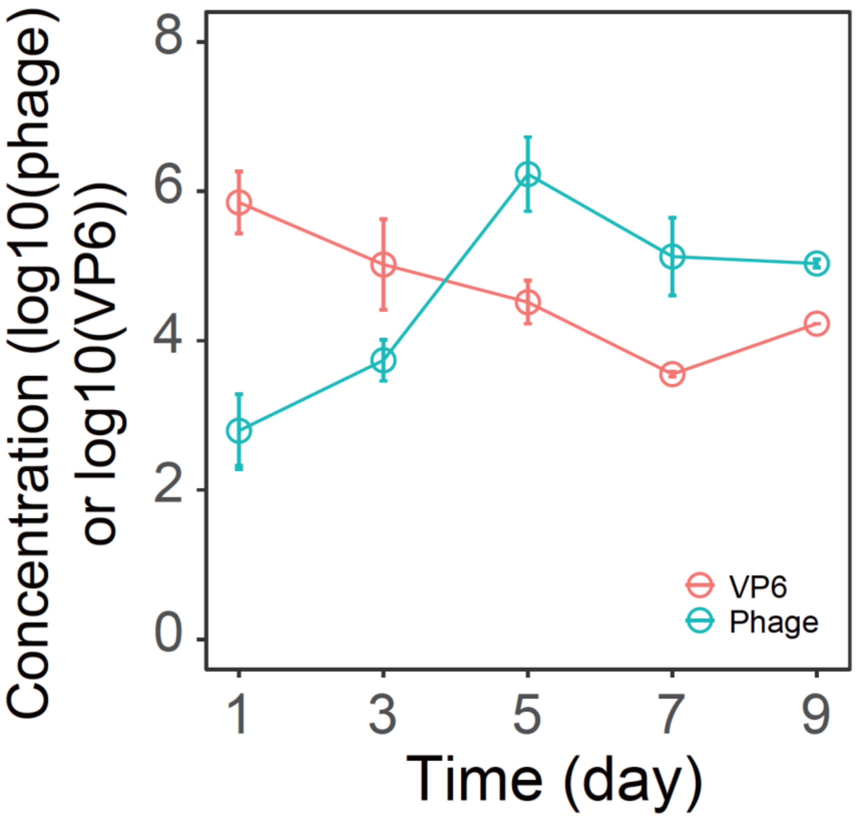


**Extended Data Fig. 8.** Dynamic changes of VP6 (CFU/g intestinal sample) and the phage (PFU/g intestinal sample) in the shrimp intestine. Data points represent the means ± SD of VP6 and the phage in the shrimp samples (n = 3) collected at different time points. Absolute concentrations in the y - axis are shown in the log10 scale.

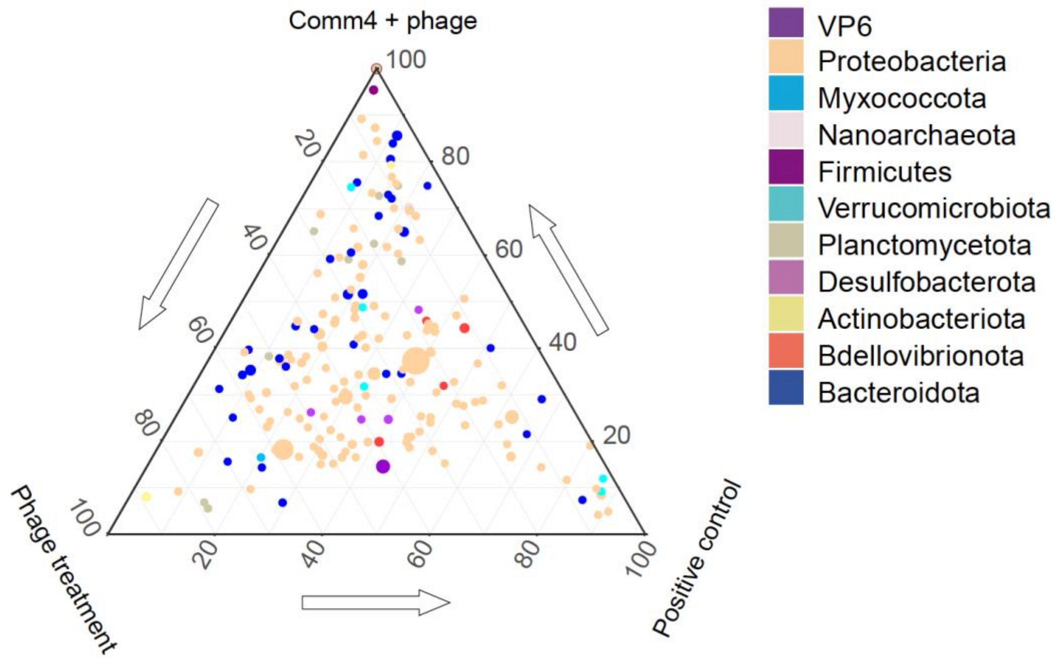


**Extended Data Fig. 9.** Ternary plot of 194 taxa in shrimp intestine to evaluate and decipher the difference effect between different treatments. Each variant represents microbial species (OTU level) and colored by phylum (except VP6). The size was proportional to the average abundance for each taxon.

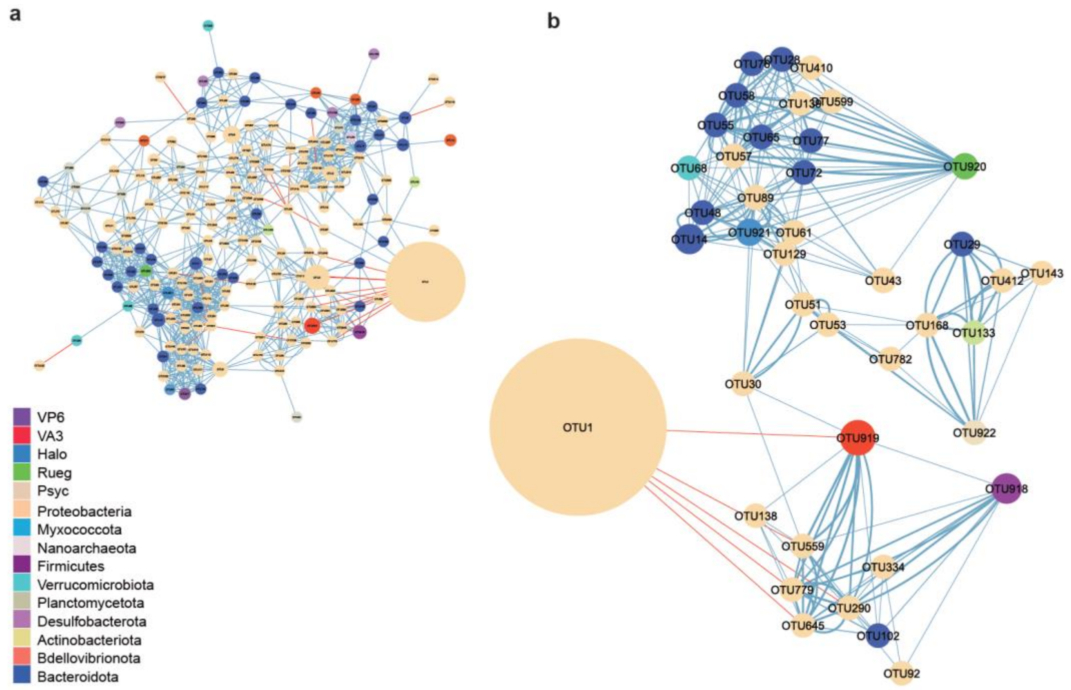


**Extended Data Fig. 10.** Co-occurrence network of Com4 and VP6 correlated with others in the shrimp intestinal microbiome. Blue and gray edges indicate significant negative and positive correlations, respectively. To highlight the important correlations, the significant level was set up with *P* value < 0.001 and R2 threshold 0.5. Nodes represent microbial OTUs (species level) and colored on the phylum level (except for the Com4 strains), with sizes proportional to the number of edges connected to the OTUs. The edge information was shown in Table S3. This network was visualized and analyzed by Cytoscape (v3.10.2)(*3*) using the edge-weighted spring-embedded mode for coinfection network.

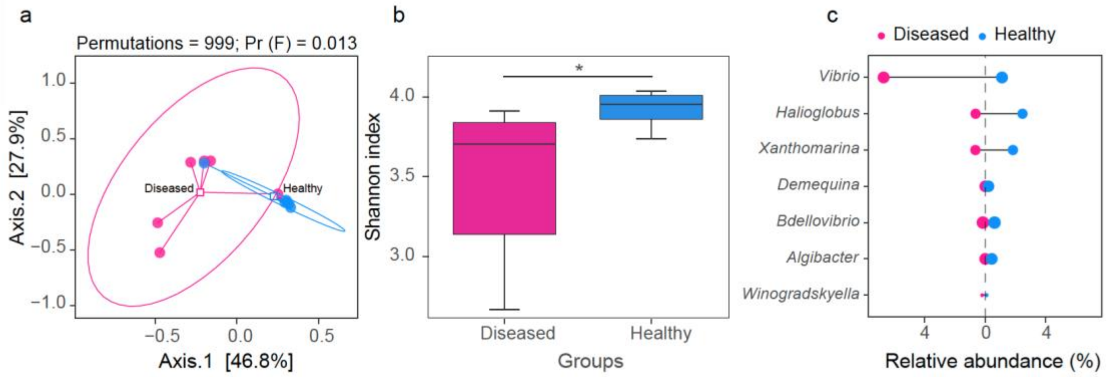


**Extended Data Fig. 11 Characterization of the intestinal microbiome of shrimp**

**a**, Between-group principal coordinate analysis (PCoA, axes 1 and 2) for the intestinal microbial composition data (16S rRNA) of shrimp. PCoA analysis was based on Bray-Curtis dissimilarity with default parameters. Sample points, from the diseased (n = 6, in pink) and healthy (n = 6, in blue) shrimp, are marked by colors according to the groups and annotated via ellipses with 95% confidence. The significant difference (*P*-values) between groups was analyzed by the Adonis function using the distance matrices and the permutation test with pseudo-*F* ratio. **b**, Alpha diversity by the Shannon index was compared using the Wilcox-test. The significant difference is donated by asterisks (*, $P < 0.05$). **c**, Discrimination of biomarkers among the top 92 genera using Random Forests with Boruta feature selection. Different colors and directions display the relative abundance of these biomarkers in diseased (pink colored) and control healthy (blue colored) shrimp. The circle scale represents the importance value (from 1.5 to 5.1) contributing to the discrimination between two groups.

**Extended Data Table 1**. Taxonomy information of 14 species of the synthetic intestinal microbiome consortia

| ID | abbreviation | phylum | class | order | family | genu | species | accession number |
|---|---|---|---|---|---|---|---|---|
| 1 | Micr | Actinobacteria | Actinobacteria | Micrococcales | Microbacteriaceae | Microbacterium spp | | SZXJY0015 |
| 2 | Exig | Firmicutes | Bacilli | Bacillales | BacillalesFamilyXIIIncertaeSedis | Exiguobacterium spp | | SZXJY0016 |
| 3 | Plan | Firmicutes | Bacilli | Bacillales | Planococcaceae | Planococcus spp | | SZXJY0017 |
| 4 | Pseu | Proteobacteria | Gammaproteobacteria | Alteromonadales | Pseudoalteromonadaceae | Pseudoalteromonas spp | | SZXJY0018 |
| 5 | Deme | Actinobacteria | Actinobacteria | Micrococcales | Demequinaceae | Demequina spp | | SZXJY0019 |
| 6 | Rueg | Proteobacteria | Alphaproteobacteria | Rhodobacterales | Rhodobacteraceae | Ruegeria spp | | SZXJY0020 |
| 7 | VA3 | Proteobacteria | Gammaproteobacteria | Vibrionales | Vibrionaceae | Vibrio spp | Vibrio alginolyticus | SZXJY0021 |
| 8 | Psyc | Proteobacteria | Gammaproteobacteria | Pseudomonadales | Moraxellaceae | Psychrobacter spp | | SZXJY0022 |
| 9 | VP6 | Proteobacteria | Gammaproteobacteria | Vibrionales | Vibrionaceae | Vibrio spp | vibrio parahaemolyticus | SZXJY0023 |
| 10 | Halo | Proteobacteria | Alphaproteobacteria | Rhodobacterales | Rhodobacteraceae | Halocynthiibacter spp | | SZXJY0024 |
| 11 | Aest | Bacteroidetes | Flavobacteriia | Flavobacteriales | Flavobacteriaceae | Gaetbulibacter spp | | SZXJY0025 |
| 12 | Shew | Proteobacteria | Gammaproteobacteria | Alteromonadales | Shewanellaceae | Shewanella spp | | SZXJY0026 |
| 13 | Tena | Bacteroidetes | Flavobacteriia | Flavobacteriales | Flavobacteriaceae | Tenacibaculum spp | | SZXJY0027 |
| 14 | Algo | Bacteroidetes | Cytophagia | Cytophagalesf | Cyclobacteriaceae | Algoriphagus spp | | SZXJY0028 |

**Extended data Table 2**. Spot tests of phages against each of the isolated commensal bacterial strains in this study. 10 µL of high dense of phage lysate (~10^10 PFU/mL) of VP6phageC was spotted onto lawns of each tested strains and incubated for 24 h or 48 h. Black/gray filling indicates whether clear zone observed or not on the bacterial lawn.

| | Strain | Infection |
|---|---|---|
| 1 | Micr | |
| 2 | Exig | |
| 3 | Plan | |
| 4 | Pseu | |
| 5 | Deme | |
| 6 | Rueg | |
| 7 | VA3 | |
| 8 | Psyc | |
| 9 | VP6 | |
| 10 | Halo | |
| 11 | Aest | |
| 12 | Shew | |
| 13 | Tena | |
| 14 | Algo | |

**Extended data Table 3**. Primer list used for qPCR assay. The prophage-specific primers (pro_VPP2) amplify a region associated with phage replication, while the internal control primers target the *gyrB* gene to normalize expression levels.

| Primer set | Direction | Sequence (5’ to 3’) |
|---|---|---|
| Pro_VPP2 | Forward | 5’-CCTCGCCATGACAAAGGACG-3’ |
| | Reverse | 5’-TGGGCTGTGATGAGCGTAGC-3’ |
| Internal control | Forward | 5’-GTGACTCTGCGGGTGGTTCA-3’ |
| | Reverse | 5’-CACGACCGATACCACAGCCA-3’ |
